# The N-terminal executioner domains of NLR immune receptors are functionally conserved across major plant lineages

**DOI:** 10.1101/2022.10.19.512840

**Authors:** Khong-Sam Chia, Jiorgos Kourelis, Martin Vickers, Toshiyuki Sakai, Sophien Kamoun, Philip Carella

## Abstract

Nucleotide-binding domain and leucine-rich repeat (NLR) proteins are a prominent class of intracellular immune receptors that are present across diverse plant lineages. However, our understanding of plant NLR structure and function is limited to the evolutionarily young flowering plant clade (angiosperms). Here, we describe an extended spectrum of NLR diversity across major plant lineages and demonstrate functional conservation of N-terminal ‘executioner’ domains that trigger immune responses. We show that broadly distributed CC (coiled-coil) and TIR (toll/interleukin-1 receptor) domains retain executioner function through trans-lineage activation of immune-related cell death in the model angiosperm *Nicotiana benthamiana*. Further examination of a CC subfamily specific to non-flowering lineages uncovered an essential N-terminal MAEPL motif with functional similarity to resistosome-forming CC-NLRs. Ectopic activation of the MAEPL-type CC in the divergent liverwort *Marchantia polymorpha* led to profound growth inhibition, defense gene activation, and signatures of cell death resembling CC activity in flowering plants. Moreover, comparative macroevolutionary transcriptomics in *Marchantia* and *Nicotiana* identified conserved CC responsive genes, providing further insight into the core aspects of CC function shared between flowering and non-flowering plants. Our findings highlight the need to understand NLR structure and function across the full spectrum of plant diversity.

## INTRODUCTION

Immune receptors play a central role in perceiving and responding to host cell invasion by parasitic organisms. In plants, decades of functional genetic research has solidified the role of nucleotide-binding domain and leucine-rich repeat (NLR) proteins as intracellular resistance (R) gene receptors that provide robust defenses against pathogen infection^1,2^. Although genomics studies have recently revealed the occurrence of NLRs across a diverse range of land plants and their algae predecessors^3,4^, our understanding of NLR function is limited to the angiosperm lineage (flowering plants). In fact, each of the ∼450 experimentally validated NLRs to date are from model or crop species of flowering plants. Given that angiosperms are a relatively young lineage, our current view of NLR diversity and evolution in plants is narrow. In particular, the extent to which plant NLRs are functionally conserved across a deep macroevolutionary timescale is unknown.

NLRs are modular proteins consisting of an N-terminal domain, a central NB-ARC domain, and a C-terminal region containing leucine rich repeats (LRR) or other superstructure forming repetitive elements^5,6^. The NB-ARC domain functions as a switch that controls the ‘on/off’ state of the receptor, whereas the variable N-terminal domain defines different NLR classes. In plants, NLR N-terminal domains include the coiled-coil (CC), G10-type CC (CC_G10_), RPW8-type CC (CC_RPW8_) and toll/interleukin-1 receptor-type (TIR) domains, whereas metazoan NLRs typically encode N-terminal PYRIN or caspase recruitment domains (CARD)^7–10^. NLR N-terminal domains are viewed as the ‘executioner’ domain encoding the biochemical activities that lead to immunity. Indeed, ectopic expression of N-terminal executioner domains in *planta*, either alone or translationally fused to fluorescent proteins like YFP, is often sufficient to activate plant immune responses^11,12^. Typical outputs downstream of NLR N-terminal domain activity include defensive hormone accumulation/signaling, transcriptional reprogramming, reactive oxygen species (ROS) accumulation, and in many cases a localized form of programmed cell death known as the hypersensitive response (HR)^13,14^.

Structural and biochemical studies have revealed the molecular functions of NLR subtypes. Upon pathogen virulence factor-dependent activation, the *Arabidopsis* CC-NLR receptor ZAR1 (hopZ1-Activated Resistance1) forms higher order oligomer complexes (‘resistosome’) in which the primary helix of each CC domain 4-helix bundle assembles into a funnel-shaped structure predicted to form pores within the plasma membrane^15,16^. In support of this idea, activated *Arabidopsis* ZAR1 or wheat Sr35 (Stem rust resistance 35) pentamers were shown to accumulate at lipid bilayers and act as non-selective cation channels *in vitro*^17,18^. This paradigm was further confirmed for the CC_RPW8_ domains of *Arabidopsis* NRG1 and ADR1, which exhibited oligomerization-dependent ion channel activity requiring the N-terminal region of the CC_RPW8_ domain^19^. By contrast, activation of the *Arabidopsis* TNL receptors RPP1 and ROQ1 induces oligomerization that reconstitutes a TIR holoenzyme complex capable of hydrolyzing NAD^+^ (nicotinamide adenine dinucleoside) to produce small molecules that bind to EDS1 (ENHANCED DISEASE SUSCEPTIBILITY1) regulatory complexes^20–23^. This, in turn, recruits CC_RPW8_-NLR receptors to execute plant immune responses^22–24^.

In plants, NLRs function at varying levels of connectivity, ranging from standalone ‘singleton’ receptors sufficient to induce NLR-mediated immunity, paired NLRs that distribute perception (sensor) and transduction (helper) activities, and in fortified networks where a minimal set of helper NLRs function alongside an extensive repertoire of sensors^25^. Within this framework, several conceptual models have emerged to explain how NLRs are poised to monitor for pathogen virulence factors (effectors) and/or their activity. While direct interactions between pathogen effectors and plant NLRs can occur, not all effectors directly interact with cognate NLR receptors^1^. In many instances NLRs monitor (or *guard*) the integrity of key host proteins and activate immunity upon effector-mediated disruption^26^. NLRs also monitor non-functional ‘decoy’ guardees that are often related to functionally relevant host targets^27^. Strikingly, such decoy domains are frequently incorporated within NLRs themselves, with the resulting ‘integrated domain’ responsible for effector recognition being genetically fused to the canonical NLR structure^6^.

Our understanding of NLR form and function is limited to angiosperms (flowering plants), which are an evolutionarily young lineage that diverged from non-flowering ancestors approximately 209 million years ago in the Upper Triassic era^28^. The first land plants evolved from freshwater charophyte algal predecessors over 500 million years ago (Cambrian-Ordovician) and diverged into several key lineages that predate the angiosperms^29^. This includes the non-vascular seed-free bryophytes (liverworts, hornworts, and mosses), vascular seed-free lineages like lycophytes (clubmosses) and monilophytes (ferns and horsetails), and the seed-bearing but non-flowering gymnosperms (conifers, cycads, ginkgos, and gnetophytes). Genomics data demonstrates that NLRs are present across land plants and in some of their algal predecessors^3,4,30^. However, functional analyses of NLRs from non-flowering plants are lacking. In particular, the extent to which NLR immune receptors or their N-terminal executioner domains are functionally conserved across the full spectrum of plant evolution is unknown.

In this study, we undertook a comparative macroevolutionary approach to understand the extent to which NLRs are functionally conserved across divergent plant lineages. Using the NLRtracker annotation tool^8^, we defined the NLR immune receptor repertoires of distantly related land plant and algal genomes for comparison against an extensive set of angiosperm NLRs^8,31^. Phylogenetic analysis of the central NB-ARC coupled with orthology analysis of N-terminal domains revealed the diversity and evolutionary history of NLR subtypes across plant lineages. *In planta* expression screening of widely distributed NLR N-terminal executioner domains (CC, CC_RPW8_, and TIR) sampled across divergent plants and algae demonstrated deep functional conservation of HR cell death induction in *N. benthamiana*. Further examination of a novel CC domain subtype enriched in non-flowering plants uncovered molecular and mechanistic similarities to angiosperm CC domains. Moreover, phenotypic dissection of non-flowering CC activity in *Nicotiana* and the model liverwort *M. polymorpha* identified shared aspects of the CC response despite over 450 million years of divergence. Collectively, our data reveal deep evolutionary conservation of NLR immune receptor executioner domains in plants and hints toward an ancestral CC-mediated immune program.

## RESULTS

### Major plant lineages harbor an extended spectrum of NLR immune receptors

To understand the diversity of NLR immune receptors present across distantly related green plants, we applied the NLR annotation tool ‘NLRtracker’^8^ to representative non-flowering plant genomes. While our primary focus was on terrestrial seed-free plants (bryophytes, lycophytes, and monilophytes), we included two streptophyte algae and a gymnosperm (seed-bearing but non-flowering) for comparison alongside an experimentally validated set of angiosperm NLRs (RefPlantNLR)^8^. Collectively, our set includes 19 organisms and captures green lineage diversity across >500 million years of evolutionary divergence, inclusive of freshwater aquatic algae (sister to all terrestrial plants), non-vascular/seed-free bryophytes, vascular but seed-free lycophytes/monilophytes, and non-flowering gymnosperms alongside angiosperm NLRs (Figure 1A). NLRtracker identified full length NLRs, degenerated NLRs, and additional NB-ARC-containing proteins (other) in all tested species (Data S1). Large NLR repertoires were observed in mosses (especially *Ceratodon purpureus*; >231), the terrestrial fern *Ceratopteris richardii* (97), and the gymnosperm *Ginkgo biloba* (248) (Figure 1B). An analysis of NB-ARC domain classification revealed an expansion of TIR-type NB-ARC domains in *Ceratodon* and *Ginkgo*, whereas the remaining non-flowering NB-ARC domains were classified as ‘Other’ or ‘undetermined’ (Figure 1C).

**Figure 1.**
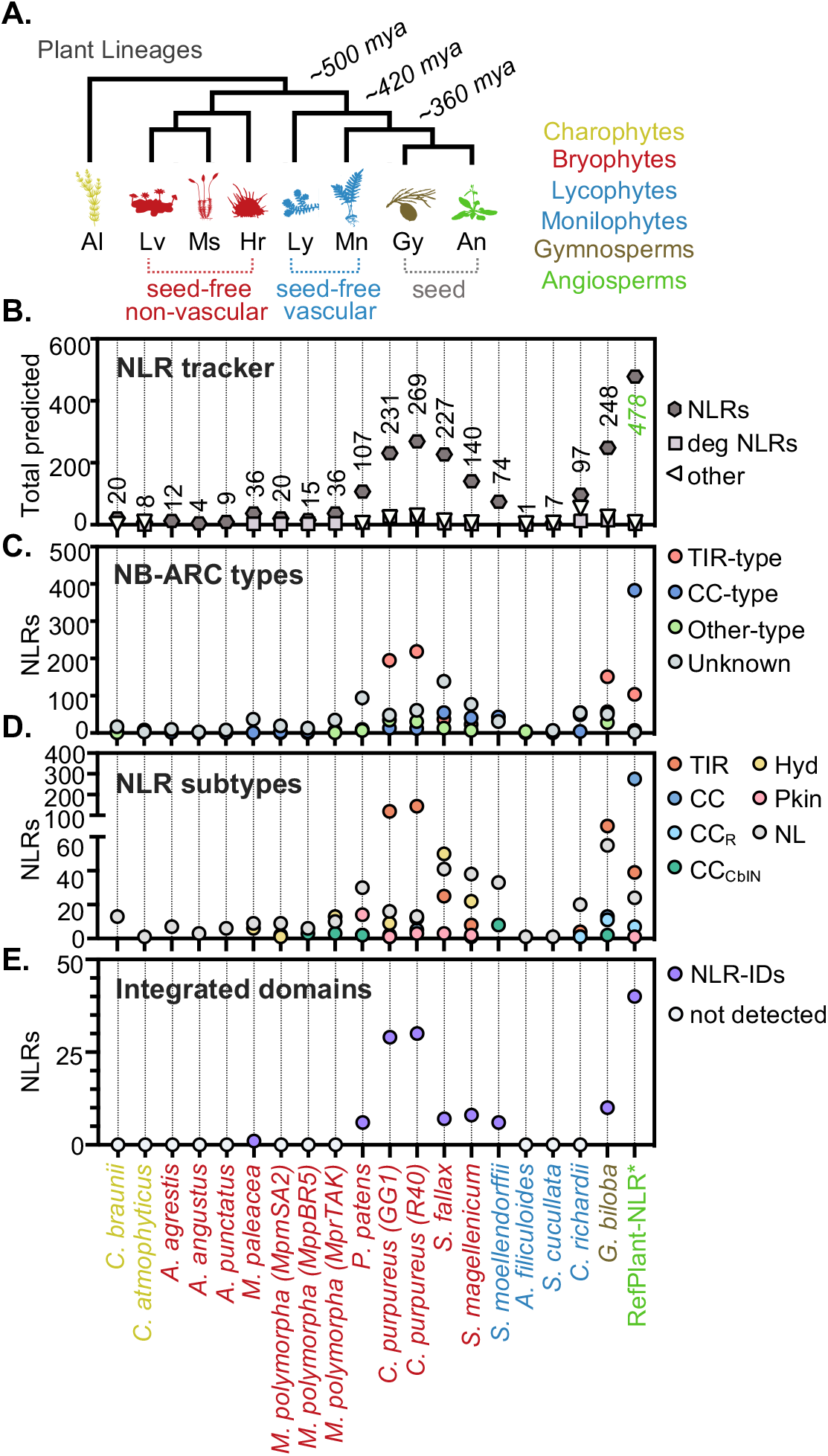
Major plant lineages harbor a diversity of NLR immune receptors. **(A)** Graphical representation of the evolutionary history of major plant lineages that includes streptophyte algae (Al), liverworts (Lv), mosses (Ms), hornworts (Hr), lycophytes (Ly), monilophytes (Mn), gymnosperms (Gy), and angiosperms (An). The indicated transitions represent a timescale of millions of years ago (mya) based on previous estimates^29^. Not to scale. **(B)** Total number of full-length NLRs (NLR), degenerated NLRs (Deg), or other NB-ARC domain-containing proteins predicted by the NLRtracker annotation tool. Numbers on the graph represent the total number of full-length NLRs predicted per species/group. **(C)** Diversity of NB-ARC domain subtypes per species/group as predicted by NLRtracker. **(D)** Diversity of NLR receptor subtypes per species/group as predicted by NLRtracker. Categories are based on predicted N-terminal executioner domains and include TIR-type (TIR), CC-type (CC), RPW8-type (CC_R_), CblN-type (CC_CblN_), hydrolase-type (Hyd), protein kinase-type (Pkn), and undefined/minimal NB-ARC-LRR type receptors (NL). **(E)** Total number of NLR immune receptor integrated domains (NLR-IDs) predicted per species/group by NLR tracker.

Further characterization of NLR subdomain composition revealed a diverse set of immune receptor architectures that included subtypes common to angiosperms (TIR, CC, and CC_RPW8_), with notable expansions of TIR-NLRs in mosses and *Ginkgo*. Moreover, NLRtracker identified receptor subtypes specific to non-flowering lineages, which included receptors containing an N-terminal αβ-hydrolase (Hyd-NLRs), protein kinase (Pkn-NLRs), or a 4-helix bundle (Cbl-N domain) with homology to the N-terminus of the human CBL (Casitas B-lineage Lymphoma) proto-oncogene (Figure 1D). We henceforth included the Cbl-N domain in the broader CC subtype classification since AlphaFold2-based structural predictions suggest a 4-helix bundle conformation reminiscent of angiosperm CC and CC_RPW8_ domains (Figure S1). Lastly, we cataloged NLRs harboring integrated domains (NLR-IDs) at the N or C-terminus of the receptor. This revealed an expansion of NLR-IDs in mosses and *Ginkgo*, whereas the remaining non-flowering lineages generally lacked integrated domains (Figure 1E, Data S1). NLR-IDs represent ∼4% of the combined non-flowering NLRome. Altogether, NLRtracker confirms the widespread prevalence of NLRs across green plants and highlights the expanded diversity of immune receptors in non-flowering lineages.

### NLRs share deep evolutionary ancestry despite N-terminal executioner domain diversity

We further explored the evolutionary history and diversification of NLRs using orthology analysis (Orthofinder) of N-terminal executioner domains alongside maximum likelihood phylogenetic analysis of the central NB-ARC (Figure 2A). Orthofinder identified 196 orthologous protein groups (orthogroups) amongst NLR N-terminal domains in green plants (Data S1). To focus on the most prevalent domains we prioritized orthogroups (OGs) within 2 or more species that had at least 10 unique loci, which revealed 11 major groups (Figure 2B). The OG0 TIR domain was the largest and most widely distributed orthogroup, followed by OG1 (αβ-hydrolase), OG2 (TIR), OG3 (Cbl-N-type CC), OG4 (protein kinase), OG6 (ZAR1-type seed plant CC), and OG15 (ADR1/NRG1-type CC_RPW8_). We also identified putative CC-like (OG7, OG8, OG13) and small/truncated beta barrel-like N-terminal domains (OG18), with structures predicted by AlphaFold2 (Figure S1). The majority of these orthogroups were enriched in representative non-flowering lineages, with exception of OG0 (widely distributed TIR), OG6 (seed plant ZAR1-type CC), and OG15 (seed plant ADR1/NRG1-type CC_RPW8_). Phylogenetic analysis of the NB-ARC domain demonstrated a clear separation of TIR and CC-type NLR clades irrespective of lineage. Similar to previous work on *M. polymorpha* and *Physcomitrium patens* NLRs^3,4^, our NB-ARC phylogeny places the vast majority of hydrolase and protein-kinase containing NLRs within a larger clade of TIR-NLRs (Figure 2C), suggesting that Hyd-NLRs and Pkn-NLRs were derived from an ancestral TIR-type NB-ARC. The only exception to this was the presence of two *C. purpureus* (moss) Pkin-NLRs within the CC clade, indicating an independent birth of these two receptors. Several atypical NLR-proteins were sister to the CC clade, including the TNP family^32^ of TIR-NBARC-TPR receptors originating in algae and present in land plants. Lastly, the highly prevalent CC-type OG3 domain enriched in non-flowering lineages was embedded within the CC clade between NLRs harboring the CC_RPW8_ and angiosperm CC domains. Collectively, these analyses reveal the deep evolutionary history of NB-ARC-containing NLR immune receptors and further highlight the diversity of N-terminal executioner domains encoded by non-flowering plants.

**Figure 2.**
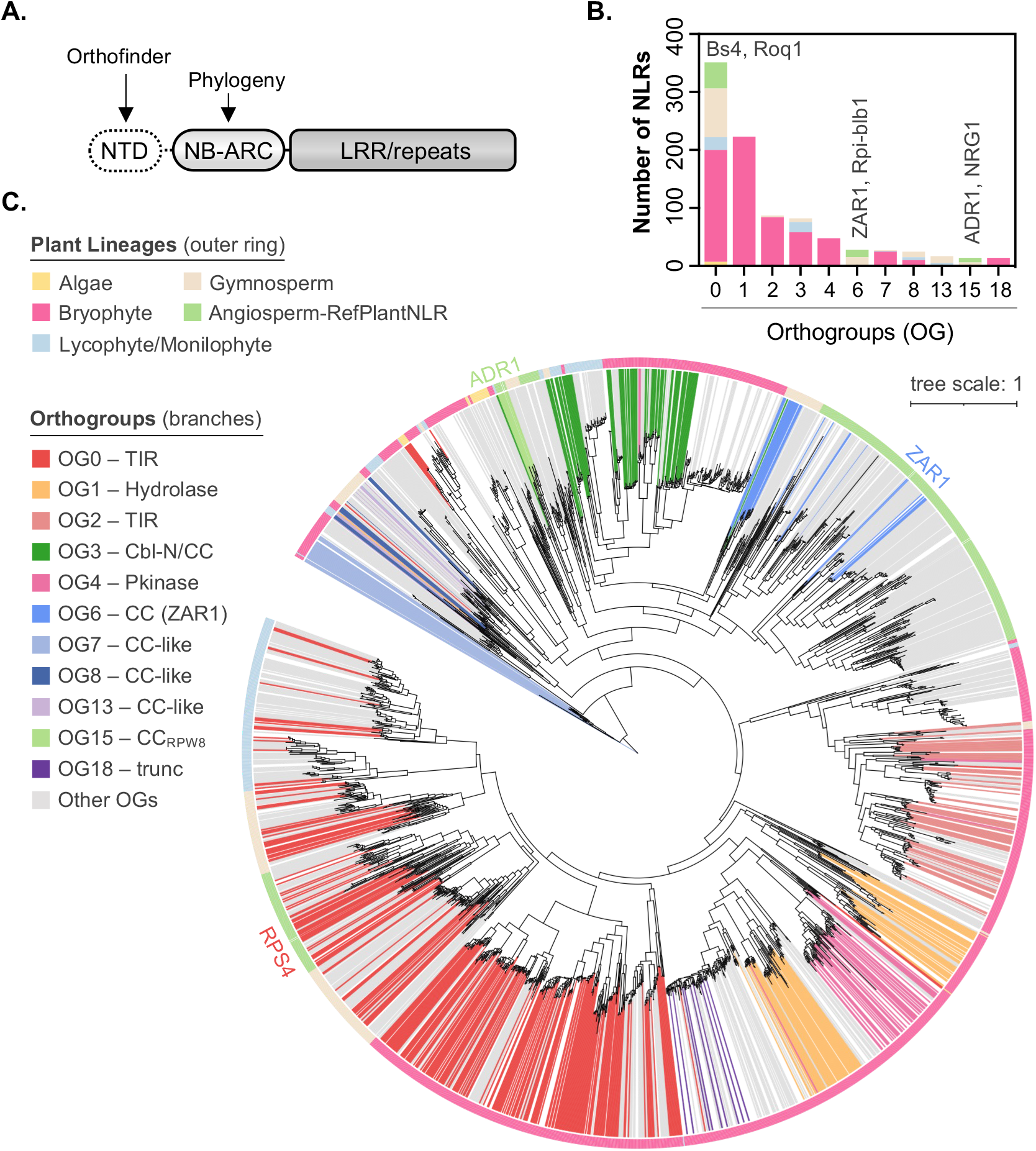
NLRs with diverse N-terminal executioner domains share deep evolutionary ancestry. **(A)** Schematic overview of canonical NLR immune receptor structure including the N-terminal executioner domain (NTD), the central NB-ARC regulatory domain, and C-terminal leucine rich repeats (LRR) or other repeats. Arrows indicate the bioinformatic analyses being conducted on each protein domain. **(B)** Frequency of key NLR N-terminal domain orthogroups (OG) observed per lineage/group. Where appropriate, representative angiosperm NLRs are listed above the respective OG. **(C)** Maximum likelihood phylogeny of diverse plant NLRs based on the central NB-ARC regulatory domain. Coloration of the outer ring represents host lineage/group while branch colors indicate key N-terminal domain OGs. A representative angiosperm TIR-NLR (RPS4), CC-NLR (ZAR1), and CC_RPW8_-NLR (ADR1) are indicated.

### The N-terminal executioner domains of NLR immune receptors are functionally conserved across major plant lineages

Our phylogenetic and orthology analyses demonstrate the ubiquitous nature of NLR subtypes across major plant lineages. However, the extent to which divergent NLRs are functionally conserved over a macroevolutionary timescale remains unknown. To address this, we screened the N-terminal executioner domains of diverse TIR and CC-type receptors for their ability to activate HR cell death when transiently expressed as eYFP fusions in the angiosperm *N. benthamiana* (Figure 3A, Data S2). We observed moderate-to-strong cell death phenotypes in 5 of 20 TIR domains cloned from non-flowering plants, with those from mosses (*P. patens* and *Sphagnum fallax*), ferns (*C. richardii*), and gymnosperms (*G. biloba*) exhibiting activity stronger than the *Arabidopsis* RPS4 control (Figure 3B; Data S2B). TIR fusions were also screened in *N. tabacum* since TIR-mediated cell death is typically stronger in this species. As anticipated, enhanced TIR-mediated cell death was observed in *N. tabacum* relative to *N. benthamiana* (Data S2C). In comparison to TIRs, a diverse collection of CC-type domains induced strong HR cell death in *N. benthamiana* (Figure 3C; Data S2D). This covered approximately 70% of tested domains from all non-flowering lineages alongside the angiosperm MLA10 (CC) and ADR1 (CC_RPW8_) controls. Predicted CC-type domains from the streptophyte alga *Chara braunii* (OG97, OG164, OG165) were used as an outgroup to land plants (Figure S1), with moderate-to-strong activity observed for 3 of 5 tested domains. In addition, we independently assayed each of the two CC domains present within predicted *M. polymorpha* tandem CC-CC-NLR receptors (Mp3g09180 and Mp3g09210), identifying HR-induction for the internal CC of Mp3g09210 only. Importantly, immunoblots confirmed stable expression of CC/TIR-eYFP fusion proteins, apart from a minority of domains inducing strong cell death phenotypes (Data S3A). Together these data demonstrate the conserved functionality of TIR and CC-type domains, highlighting a deep evolutionary origin of NLR immune receptor executioner domains in plants.

**Figure 3.**
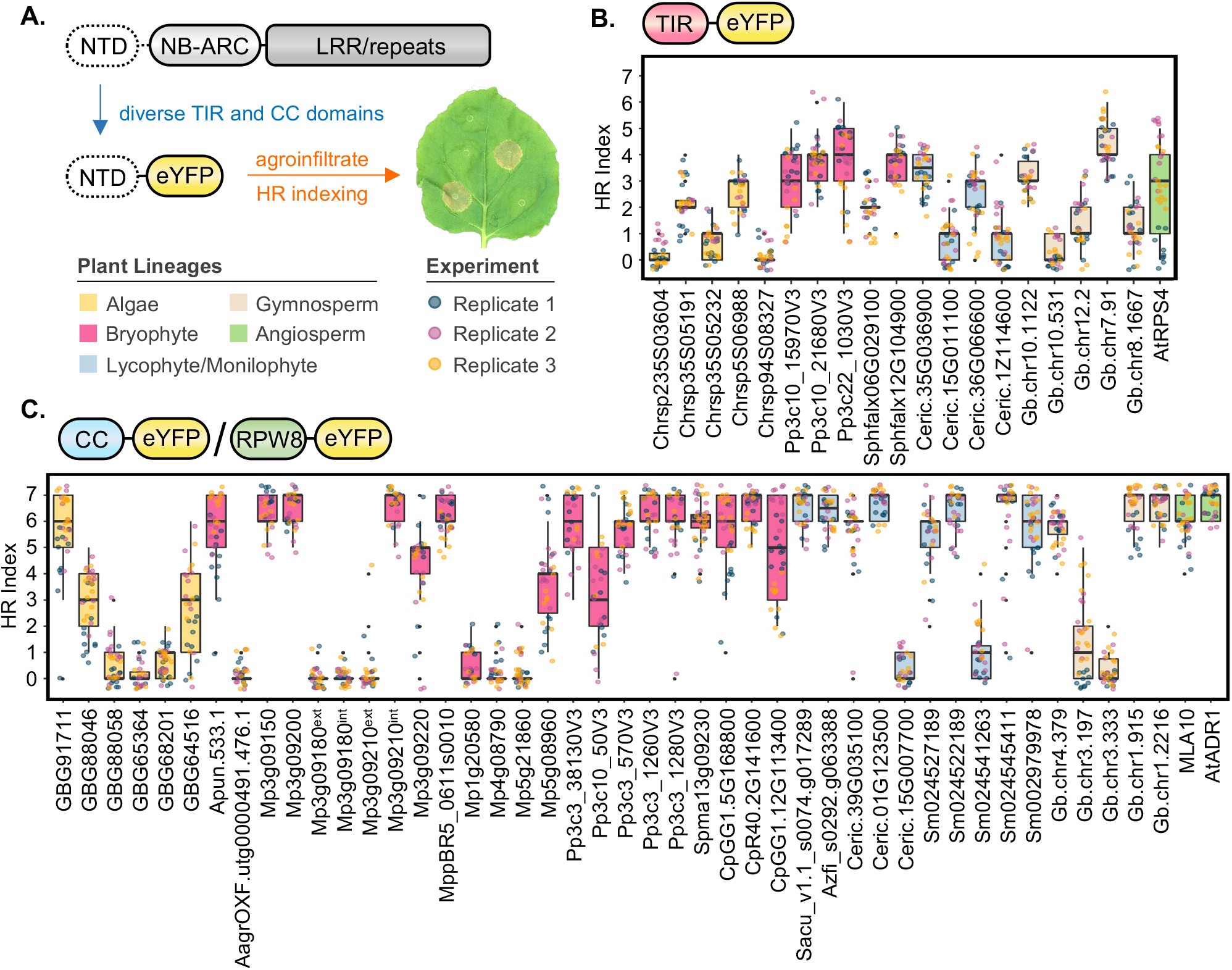
Widely distributed N-terminal executioner domains are functionally conserved across major plant lineages. **(A)** Schematic overview of the experimental design, whereby diverse TIR and CC domains are fused to eYFP, transiently expressed in *Nicotiana*, and scored for their ability to induce immune related hypersensitive response (HR) cell death via the HR index (from 0-7). Examples of macroscopic cell death phenotypes in an *N. benthamiana* leaf are depicted. **(B)** HR cell death induction of TIR-eYFP fusions transiently expressed in *N. benthamiana* leaves. Scoring (HR index) was performed 5 days post agroinfiltration, Data from for three independent experimental replicates are shown (n ≥ 9 infiltrations per replicate). **(C)** HR cell death induction of CC/RPW8-eYFP fusions transiently expressed in *N. benthamiana* leaves. Scoring (HR index) was performed 5 days post agroinfiltration. Data from three independent experimental replicates are shown (n ≥ 9 infiltrations per replicate).

### Non-flowering plants encode a unique CC_OG3_ domain that harbors a sequence conserved N-terminal ‘MAEPL’ motif

The broad distribution of functionally conserved CCs suggests that common principles underpin their activity despite over 500 million years of green plant evolution. To identify conserved and functionally relevant features in divergent CCs, we queried the highly prevalent CC_OG3_ domains of non-flowering plants for enriched amino acid sequence motifs using MEME (Multiple EM for Motif Elicitation)^33^. While this revealed several conserved motifs across the entire CC_OG3_ domain (Data S4AB), we focused on two adjacent motifs present at the N-terminus given their prevalence (79/80 and 38/80) and because similarly situated motifs underpin CC function in angiosperms^19,34^. Sequence alignment of this region revealed a conserved consensus motif at the CC_OG3_ N-terminus that we named “MAEPL” (Figure 4A; Data S4C). We used this alignment to build a Hidden Markov Model (HMM) profile for further examination of MAEPL prevalence across green plant proteomes via the HMMER tool^35^. Overall, HMM profiling of non-flowering proteomes identified high-scoring MAEPL matches in NLRs predicted by NLRtracker (Figure S2A). In comparison, an analysis of MAEPL prevalence in the well-annotated angiosperm proteomes of *A. thaliana* and *Solanum lycopersicum* identified only low scoring hits in NLR and non-NLR proteins alike (Figure S2B). To extend this comparison further, we interrogated the angiosperm NLR atlas (>90,000 NLRs)^31^ and our combined set of non-flowering lineage NLRs for the presence MAEPL-like motifs, which revealed an increased occurrence of high-scoring motifs in non-flowering plants relative to angiosperms (Figure 4B; Figure S2C). In contrast, the similarly situated N-terminal MADA^34^ motif of angiosperm CC domains was enriched in angiosperms but not non-flowering plants (Figure 4B; Figure S2D). Despite these differences, alignments comparing non-flowering MAEPL and angiosperm MADA N-terminal motifs demonstrate intriguing similarities in their composition (Figure 4C; Data S4D), including commonly situated leucine residues critical for MADA function in angiosperms^34^. Taken together, these results hint to a common framework for CC N-terminal motif functionality that diverged as angiosperms split from non-flowering predecessors.

**Figure 4.**
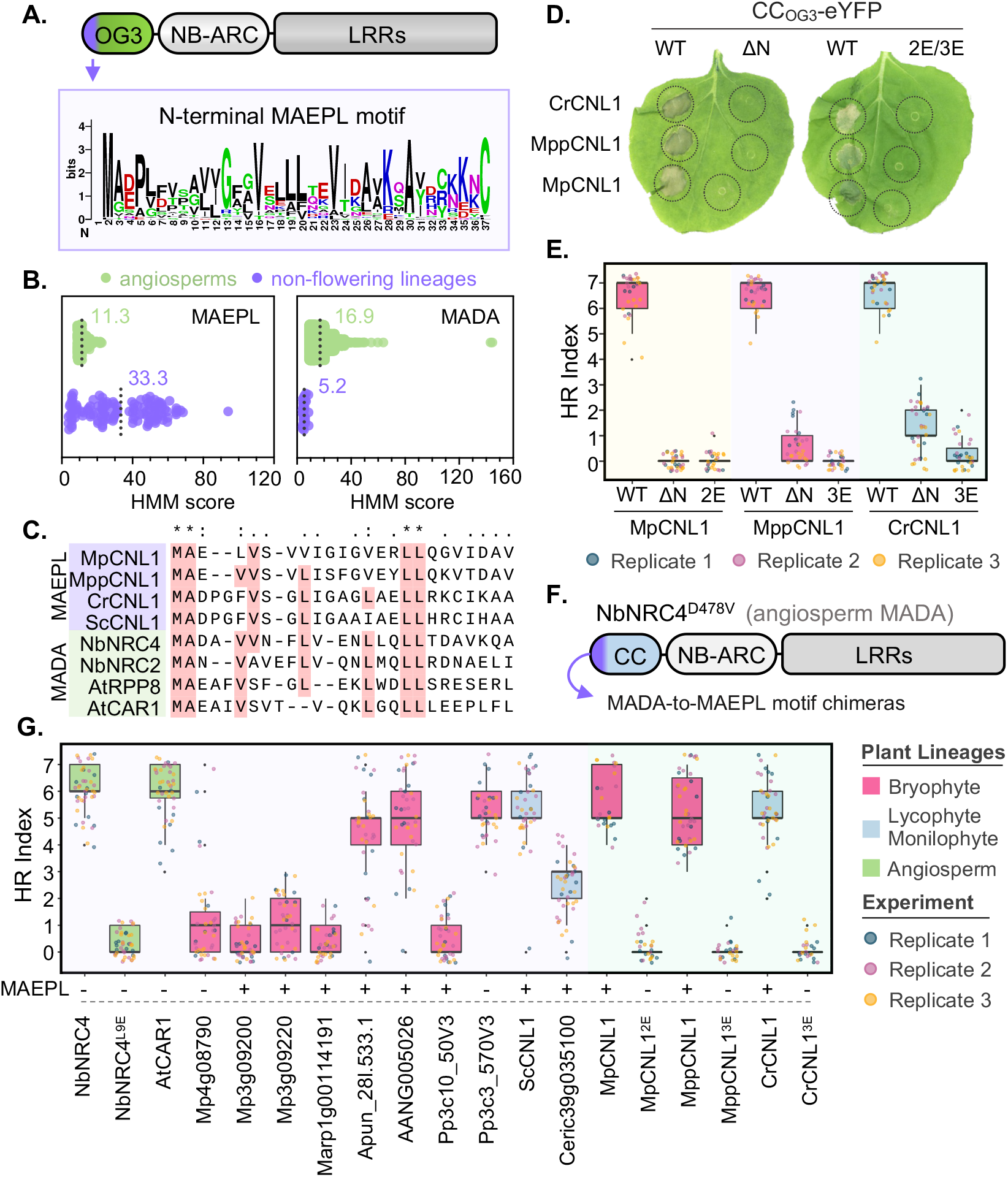
The sequence conserved MAEPL motif is essential for CC_OG3_ domain activity and is functionally analogous to the angiosperm MADA motif. **(A)** Schematic representation of a CC_OG3_-NLR immune receptor. The location of the MAEPL motif on the CC domain is indicated by an arrow and the consensus amino acid sequence of the motif is illustrated using WebLogo (https://weblogo.berkeley.edu/logo.cgi). **(B)** Hidden Markov model (HMM) profiling of the N-terminal MAEPL and MADA motifs in non-flowering NLR immune receptors identified in this study (non-flowering) relative to the angiosperm NLR atlas^31^ (angiosperms). Mean motif scores are indicated on each graph by a numerical value and a dotted line. **(C)** Amino acid sequence alignment of MAEPL and MADA motifs in representative CC domains. Conserved residues are indicated by an asterisk (*) above the alignment, similar residues by dots. For non-flowering NLRs, gene symbols correspond to MpCNL1 (Mp3g09150), MppCNL1 (MppBR5_0611s0010.1), CrCNL1 (Ceric.01G123500.1.p), and ScCNL1 (Sacu_v1.1_s0074.g017289). **(D)** Macroscopic HR cell death phenotypes of CC_OG3_-eYFP fusions comparing wild-type domains (WT), N-terminal MAEPL truncations (ΔN), and L-to-E MAEPL variants (MpCNL1^L16/17E^/2E; MppCNL1^L8/16/17E^/3E; CrCNL1^L10/18/19E^/3E) transiently expressed in *N. benthamiana*. Images were obtained 5 days post agroinfiltration and are representative of 3 independent experiments. **(E)** Quantification of HR cell death caused by CC_OG3_-eYFP (WT), N-terminal trunctions (ΔN), and L-to-E variants (2E/3E) for MpCNL1, MppCNL1, and CrCNL1 domains. Cell death was scored (HR index) 5 days post agroinfiltration. Data from three independent experimental replicates are shown (n ≥ 9 infiltrations per replicate). **(F)** Graphical representation of the MADA-to-MAEPL N-terminal motif swapping experimental design. An autoactive variant of MADA-CC-NLR NbNRC4^D478^ is used as a scaffold to assess N-terminal motif competency in *N. benthamiana*. **(G)** HR cell death induction of NbNRC4^D478V^-6HA chimeras transiently expressed in *N. benthamiana*. N-terminal motif chimeras were generated using motifs belonging to the indicated receptors. The presence of a MAEPL motif is indicated (+/-). Cell death was scored (HR index) 5 days post agroinfiltration. Data from three independent experimental replicates are shown (n ≥ 9 infiltrations per replicate).

### MAEPL is required for CC_OG3_ activity and is a functional analog of the angiosperm MADA motif

To determine the functional relevance of MAEPL we generated N-terminal truncations of three CC_OG3_-eYFP fusions (MpCNL1, MppCNL1, and CrCNL1) and assessed their ability to activate HR cell death in *N. benthamiana*. In each instance, MAEPL-truncations (ΔN) failed to induce cell death whereas wildtype MAEPL-containing CC-eYFP fusions (WT) were fully competent (Figure 4DE; Data S2E). Next, we generated MAEPL motif variants in which conserved leucine residues (hydrophobic) were replaced by glutamic acid (hydrophilic), a strategy shown to impact angiosperm MADA functionality^34^. Again, wild-type CC_OG3_-eYFP controls were fully competent while double (2E) or triple (3E) L-to-E mutations abolished HR-cell death (Figure 4DE; Data S2F). Together, these data demonstrate that the MAEPL motif is essential for CC_OG3_ function in a manner analogous to the angiosperm MADA motif. To address this similarity, we tested whether MAEPL motifs from non-flowering CCs can functionally replace MADA in an auto-active variant of the NbNRC4 angiosperm helper CC-NLR (Figure 4F). The autoactivated NbNRC4^D478V^-6HA receptor retaining its original MADA motif elicited strong HR cell death, while the MADA disrupted L9E variant was non-functional (Figure 4G; Data S2G). As a control, we generated a MADA-to-MADA chimera with the N-terminal motif of Arabidopsis AtCAR1, which effectively rescued HR cell death. Several MAEPL-NbNRC4^D478V^-6HA chimeras exhibited strong HR cell death comparable to NbNRC4^D478V^-6HA (Figure 4G; Data S2G), indicating that MAEPL can indeed replace MADA. Importantly, conserved leucine residues within the MAEPL motif were essential for this activity, as the L-to-E variants of MpCNL1, MppCNL1, and CrCNL1 chimeras failed to elicit an HR (Figure 4G; Data S2G). Immunoblots confirmed stable expression for all domains, variants, and chimeras in *N. benthamiana* (Data S3BC).

Emerging data indicates that resistosome-forming MADA-CC-NLRs associate with the plasma membrane^34,36^, which is often observed as discontinuous fluorescent punctae along the membrane in confocal fluorescence microscopy experiments. Similar to angiosperm MADA-type NLRs, the MAEPL MpCNL1 (Mp3g09150) CC_OG3_-eYFP fusion revealed clear puncta formation that discontinuously localized with a Remorin (REM1.3)-RFP plasma membrane marker, while a GUS-YFP control was nucleocytoplasmic (Figure S3A). Puncta formation was not altered in MpCNL1 2E or 3E MAEPL variants (Figure S3A), consistent with observations of stabilized MADA mutant localization in angiosperms^34,36,37^. Collectively, these data demonstrate that the divergent MAEPL motif of non-flowering land plants is functionally analogous to the angiosperm MADA.

### MAEPL-CC_OG3_ activates immune-like responses in the liverwort *Marchantia polymorpha*

The functional conservation of divergent executioner domains in *N. benthamiana* prompted us to examine whether MAEPL-CC_OG3_ activates immune responses in non-flowering plants. To address this, we used the model experimental liverwort *M. polymorpha*, a bryophyte species that diverged from *N. benthamiana* over 450 million years ago^38^. Using an estradiol-based (XVE) conditional expression system^39^, we interrogated the function of the MpCNL1 (Mp3g01950) MAEPL-CC_OG3_ domain in the wild-type TAK1 accession. The importance of the MAEPL motif was assessed by comparing the activity of full length MpCNL1^CC^-eYFP (MpC1) against an N-terminally truncated MpCNL1^CCΔN^-eYFP variant (MpC1ΔN) and an mCitrine-HA (mCit-HA) control. Severe growth inhibition was specifically observed in MpC1 liverworts cultivated directly on estrogen-supplemented media, which displayed dark pigmentation characteristic of biotic^40^ and abiotic^41^ stress in *Marchantia* (Figure 5A; Figure S4A). By contrast, MpC1ΔN and mCit-HA lines cultivated with estradiol remained healthy, similar to all DMSO controls. These results were further confirmed by exogenous estradiol application in 3 week-old liverworts, with MpC1 lines exhibiting tissue darkening and brown phenolic-like deposits at apical notches. Again, DMSO controls as well as estrogen-treated MpC1ΔN and mCit-HA remained healthy (Figure 5B; Figure S4B). Importantly, immunoblotting confirmed estradiol-dependent accumulation of the mCitrine-HA control and MpCNL1^CC^-eYFP (Data S3D). In comparison, MpC1ΔN lines exhibited reduced stability in liverwort cells such that full length MpCNL1^CCΔN^-eYFP fusions were detected alongside eYFP cleavage products (Data S3D).

**Figure 5.**
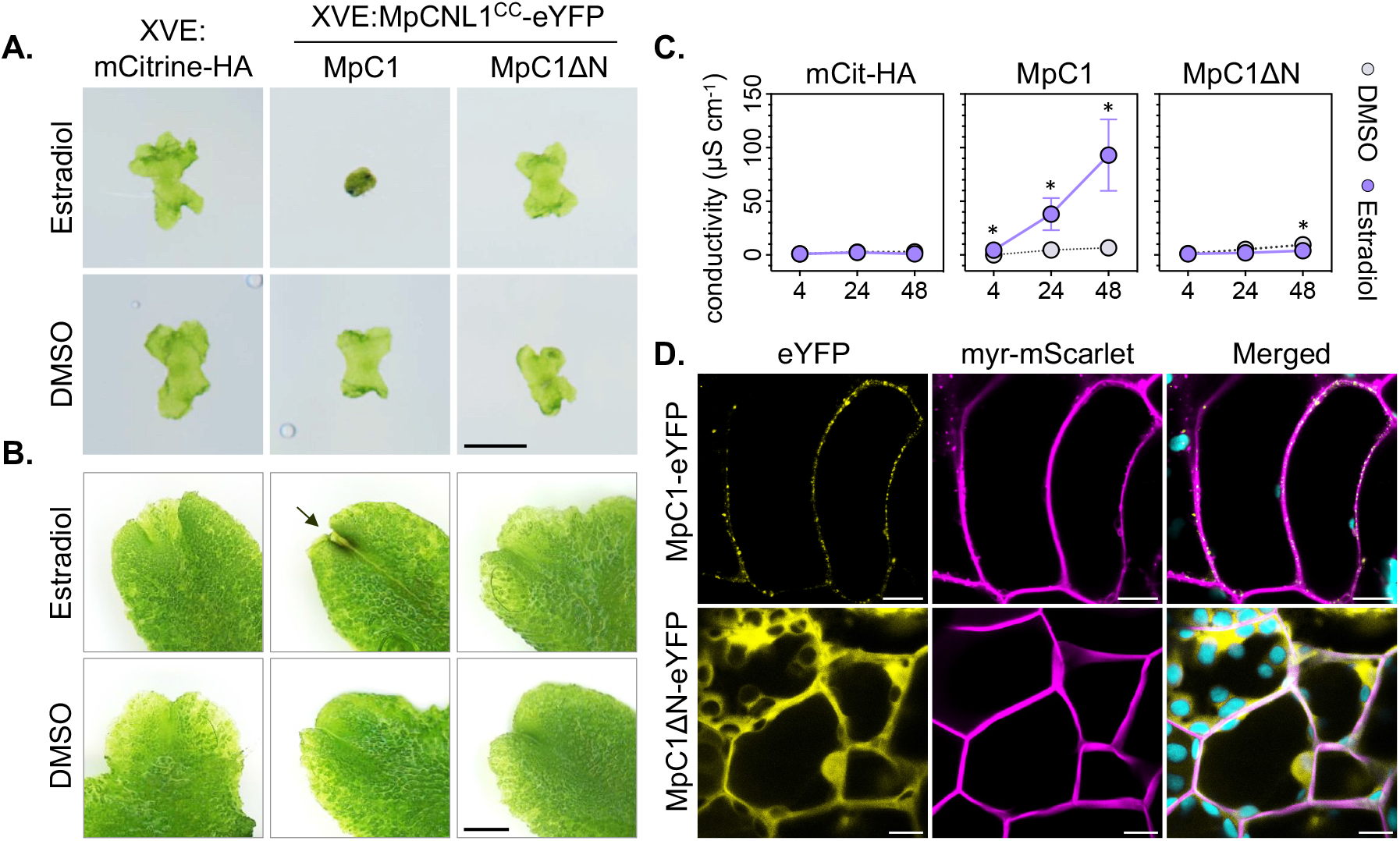
MAEPL-CC_OG3_ activates an immune-like response in the liverwort *Marchantia polymorpha*. **(A)** Macroscopic phenotypes of *Marchantia* transgenic lines XVE:mCitrine-HA (mCit-HA), XVE:MpCNL1^CC^-eYFP (MpC1, line 1), or the N-terminally truncated XVE:MpCNL1^CCΔN^-eYFP (MpC1ΔN, line 3) grown on estradiol (20 μM) or DMSO (0.1%) control media. Images are representative of growth phenotypes observed in 3 experimental replicates (n= 8 plants) at 4 days post plating. Scale bar = 2 mm. **(B)** Macroscopic phenotypes of *Marchantia* transgenic lines XVE:mCitrine-HA (mCit-HA), XVE:MpCNL1^CC^-eYFP (MpC1, line 1), or XVE:MpCNL1^CCΔN^-eYFP (MpC1ΔN, line 3) 1 day post vacuum infiltration with estradiol (50 μM) or DMSO (0.25% in water). Images are representative of phenotypes observed in 3 or more experimental replicates (n ≥ 8 plants). An arrow indicates tissue darkening at the apical notch of MpC1 liverworts. Scale bar = 2 mm. **(C)** Conductivity (μS cm^-1^) of *Marchantia* thalli treated with estradiol (50 μM) or DMSO (0.25%) at 4, 24, and 48 hours post infiltration (hpi). Statistically significant differences are denoted by an asterisk (* p< 0.05, Student’s t-test). Error bars represent standard deviation of the mean. Data from three independent experimental replicates is presented (n=12 plants per experiment). **(D)** Confocal fluorescence microscopy shows the localization of MpC1-eYFP and MpC1ΔN-eYFP alongside a myristolated-mScarlet (myr-mScarlet) membrane marker in *Marchantia polymorpha*. Images were acquired 24 hours post estradiol treatment (20 µM) in MpC1-eYFP/myr-mScarlet (XVE:MpCNL1^CC^-eYFP/MpEF1a:myr-mScarlet) and MpC1ΔN-eYFP/myr-mScarlet(XVE:MpCNL1^CCΔN^-eYFP/MpEF1a:myr-mScarlet) transgenic lines. Plastid autofluorescence is false-colored in cyan. Scale bars = 10 µm. Images are representative of 3 experimental replicates.

Next, we performed the commonly used technique of trypan blue staining^42^ to determine whether MAEPL-CC_OG3_ causes cell death in liverworts. As outlined in the methods and supporting data, we were unable to obtain conclusive results due to technical difficulties in our liverwort expression system (Fig. S4CD). We therefore performed ion leakage assays, which are routinely used to monitor membrane integrity and serve as a proxy for NLR-mediated cell death in angiosperms^43^. Here, we observed consistent increases in sample conductivity caused by ion leakage in estradiol-treated MpC1 tissues from 4-48 hpi, whereas MpC1ΔN and mCit-HA liverworts exposed to estradiol displayed minimal conductivity comparable to DMSO-treated controls (Figure 5C; Figure S4E). Collectively, these results demonstrate that the MAEPL-CC_OG3_ domain activates an immune-like response that overlaps with well-established executioner domain outputs of flowering plants.

### The MAEPL-CC_OG3_ domain forms membrane-localized puncta in liverwort cells

Since MAEPL-CC_OG3_-eYFP was observed at membrane-localized puncta in angiosperms, we performed confocal microscopy to examine whether puncta similarly form in non-flowering plants. Confocal microscopy revealed that the MpCNL1^CC^-eYFP fusion formed discontinous puncta that overlapped with plasma-membrane localized myristolated-mScarlet in liverwort cells (Figure 5D). We also observed MpCNL1^CC^-eYFP signals within intracellular inclusion bodies (Figure S3B). By comparison, N-terminally truncated MpCNL1^CCΔN^-eYFP exhibited nucleocytoplasmic distribution. Together, these data demonstrate that membrane-localized puncta formation is conserved in non-flowering plants and suggests that MAEPL-CC_OG3_ targets the membrane for immune-related activity in plant cells.

### MAEPL-CC_OG3_ activates liverwort defense gene expression

To gain a more comprehensive understanding of how the MAEPL-CC_OG3_ domain activates immune-related responses in liverworts, we performed RNA-sequencing (RNA-seq) experiments comparing gene expression profiles of estradiol-treated MpC1 (XVE:MpCNL1^CC^-eYFP line 1), MpC1ΔN (XVE:MpCNL1^CCΔN^ -eYFP line 3), and mCit-HA (XVE:mCitrine-HA) liverworts at 24 hpi. Transcriptomes of mCit-HA and MpC1ΔN were largely overlapping, whereas MpC1 liverworts displayed a drastic change in their expression profile (Figure 6A; Figure S5AB). Differential expression analysis (log2 fold change [|LFC|] ≥ 2 and adjusted p < 10^−3^) comparing MpC1 or MpC1ΔN to the mCit-HA control identified a large set of differentially expressed genes (DEGs) in MpC1 relative to the MAEPL-truncated MpC1ΔN (Figure 6B; Figure S5C; Data S5). To further support these data, we validated a subset of CC_OG3_-responsive genes by qRT-PCR (Figure S5D). Functional enrichment analyses comparing up and downregulated genes of estradiol-treated MpC1 further defined the CC_OG3_ response of liverworts. Terms associated with plant defense responses and biosynthetic activity were enriched in upregulated genes, whereas several terms associated with growth, cellular homeostatic functions, and redox activity were enriched in downregulated genes (Data S5). In comparison, only 18 transcripts were differentially expressed in MpC1ΔN versus mCit-HA controls. While this may be explained by reduced functionality and stability of MpC1ΔN-eYFP fusions, several CC_OG3_-NLRs were specifically upregulated in MpC1ΔN but not MpC1 liverworts (Data S5), which hints towards compensatory feedback caused by CC dysfunction. Altogether, RNA-seq analysis demonstrates that MAEPL-CC_OG3_ activity induces a strong molecular response that prioritizes plant defenses over normal growth and cellular functions in liverworts. This growth-defense tradeoff is reminiscent of known NLR autoactivity phenotypes in angiosperm models like *Arabidopsis*^44^, however the extent to which NLR-mediated responses are conserved across distantly related lineages remains unknown.

**Figure 6.**
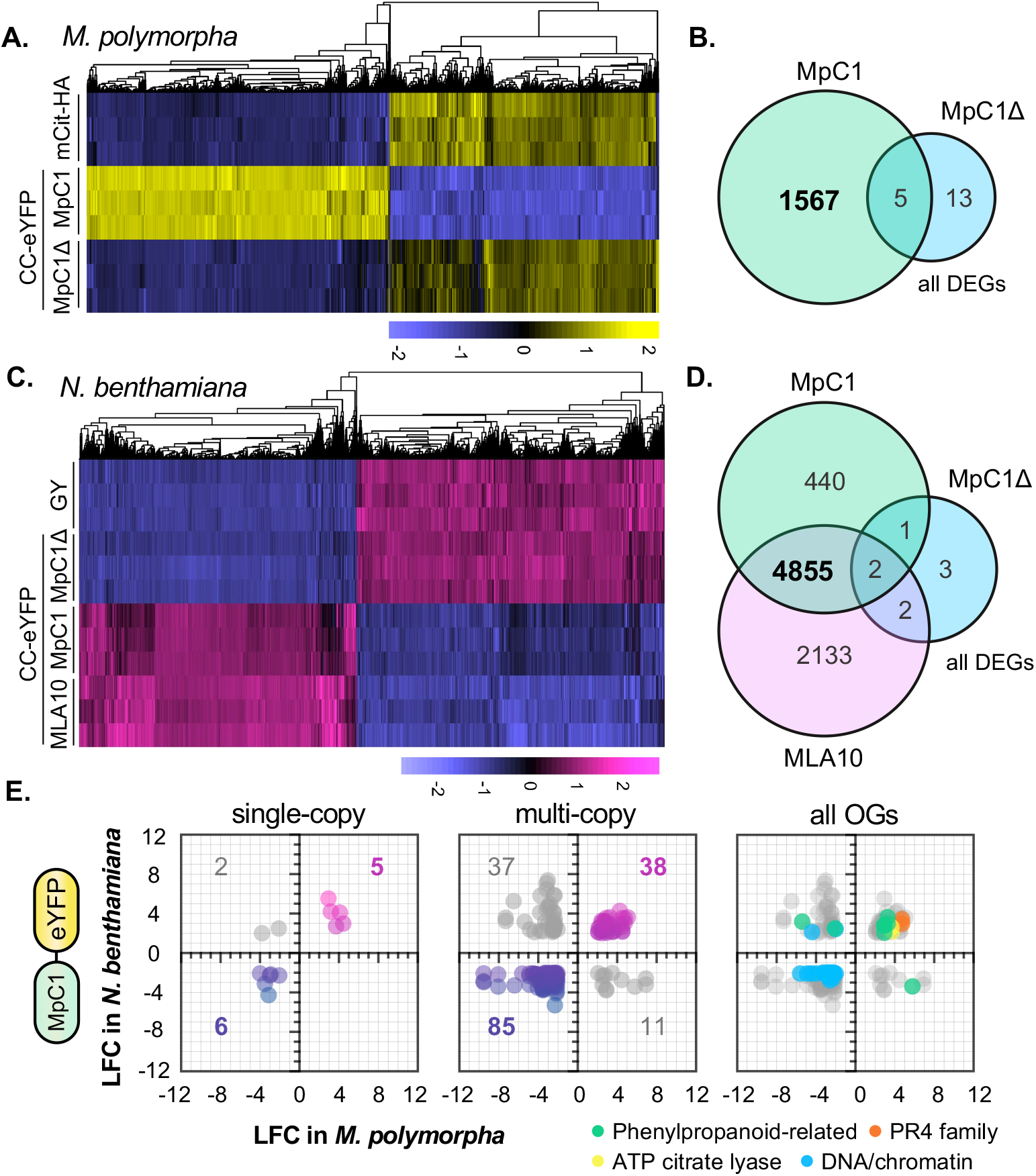
CC_OG3_ activates conserved immune-like transcriptional responses in flowering and non-flowering plants. **(A)** Hierarchical clustering of significantly differentially expressed genes in mCit-HA (XVE:mCitrine-HA), MpC1 (XVE:MpCNL1^CC^-eYFP, line 1), and MpC1Δ (XVE:MpCNL1^CCΔN^-eYFP, line 3) *M. polymorpha* transgenics 24 hours after vacuum infiltration with 20 μM estradiol (adjusted p-value < 10^−3^, log fold change (|LFC| ⩾2). Variance-stabilized row-centered counts are shown. **(B)** Total number of differentially expressed genes (DEGs) shared between *M. polymorpha* MpC1 and MpC1Δ transgenic lines. Differential expression is based on comparisons to the mCit-HA control. **(C)** Hierarchical clustering of significantly differentially expressed genes in *N. benthamiana* leaves transiently expressing GY (GUS-YFP), MpC1 (MpCNL1^CC^-eYFP), MpC1Δ (MpCNL1^CCΔN^-eYFP), or the angiosperm CC domain of MLA10 (MLA10^CC^-eYFP) at 24 hours post agroinfiltration (adjusted p-value < 10^−3^, log fold change (|LFC| ⩾2). Variance-stabilized row-centered counts are shown. **(D)** Total number of differentially expressed genes (DEGs) shared in *N. benthamiana* leaves transiently expressing MLA10, MpC1, or MpC1Δ. Differential expression is based on comparisons to the GUS-YFP control treatment. **(E)** Orthology analysis of *Marchantia* and *Nicotiana* MpCNL1^CC^-eYFP transcriptomes. Orthologous genes belonging to single or multi-copy orthogroups having an adjusted p-value < 10^−3^ and log fold change (LFC) ≥ 2 or ≤ -2 were considered. Numbers of DEGs per sector and functional enrichment categories are indicated.

### MAEPL-CC_OG3_ activates conserved immune-like transcriptional responses in flowering and non-flowering plants

The MAEPL-CC_OG3_ executioner domain activates immune-like responses in flowering and non-flowering plant species that diverged over 450 million years ago. To understand the extent to which these responses overlap, we compared the MpCNL1^CC^-eYFP activated transcriptome of *Marchantia* to the model angiosperm *N. benthamiana*. To accomplish this, we performed further RNA-seq analyses in *N. benthamiana* leaves transiently expressing MpCNL1^CC^-eYFP (MpC1), MAEPL-truncated MpCNL1^CCΔN^-eYFP (MpC1ΔN), the angiosperm MLA10^CC^-eYFP (MLA10), and a GUS-YFP control. Differential gene expression analyses revealed strong transcriptional shifts for MpC1 and MLA10 relative to the inactive GUS-YFP and MpC1ΔN controls (Figure 6C; Figure S5EF; Data S5). The expression profiles of MLA10 and MpC1 largely overlapped, with approximately 75% of all CC-upregulated genes shared between treatments (Figure 6C; Figure S5G). This suggests that the divergent CC_OG3_ behaves similar to the angiosperm MLA10 CC domain, consistent with the fact that each domain causes significant cell death in *N. benthamiana* leaves.

Next, we assessed the extent to which MpCNL1^CC^-eYFP transcriptional responses are shared between *Nicotiana* and *Marchantia*. First, we defined sets of orthologous protein-coding genes (orthogroups) using OrthoFinder^45^. This revealed a total of 6684 orthogroups, of which 1511 were single copy (one-to-one corresponding) orthologs. We observed relatively few single copy orthologs with shared expression profiles in our early timepoint CC-induction transcriptomes, with 5 commonly upregulated and 8 commonly downregulated genes (Figure 6E; Data S5). We therefore expanded our analysis to multi-copy orthologs, which revealed a larger number of genes shared between *Nicotiana* and *Marchantia*. Overall, expression profiles were generally congruent as ∼73% of orthologous DEGs were commonly up- or down-regulated (Figure 6E). Intriguingly, commonly downregulated orthologs represented the largest shared gene set between CC-induced *Marchantia* and *Nicotiana* tissues. Functional enrichment analyses identified a suite of DNA-associated machinery, including general transcription factors, replication machinery, and chromatin maintenance genes (Figure 6E; Data S5). By contrast, common CC upregulated genes included a collection of phenylpropanoid-related enzymes, ATP-citrate lyases, and members of the PR4 family. Collectively, these data suggest an evolutionarily conserved molecular response to CC activity that is likely centered on the induction of biochemical immunity alongside the repression of DNA/chromatin homeostasis.

## DISCUSSION

In this study, we demonstrate that NLR immune receptors have retained functionality in their N-terminal domains across 500 million years of plant evolution. We used the angiosperm model *N. benthamiana* to show that CC and TIR domains from algae to gymnosperms are capable of trans-lineage immune activation. Given the large evolutionary distances separating these lineages within the plant kingdom, we hypothesize that CC and TIR executioner domains arose early during plant evolution and have retained their biochemical functions throughout the conquest of land.

We functionally validated the immune activity of a diverse set of CC-type N-terminal domains across plant and algal genomes. The capacity of diverse CC domains to activate HR cell death in *N. benthamiana* reveals their ubiquitous role as executioners of plant programmed cell death. The trans-lineage activity of OG97 CC-like domains from the streptophyte alga *C. braunii* particularly supports this, as it diverged from land plants over 550-750 million years ago^29^. Fungal and metazoan N-terminal 4 helix bundle domains cause cell death in an analogous fashion to plant CC domains^12,46–48^. Intriguingly, CC domain mechanisms may be widely transferable between kingdoms as cell death was reported in metazoan cells expressing angiosperm CC-NLRs^18,19^. While this remains to be explored in further detail, these findings suggest that fundamental biochemical mechanisms may underpin the function of CC-like 4 helix bundles across the tree of life.

The CC_OG3_ domain, which generally carries the MAEPL motif at its very N-terminus, is the most common CC subtype in non-flowering lineages. We found that the MAEPL motif is essential for cell death activity in a manner analogous to the MADA motif that occurs in about 20% of angiosperm CC-NLRs^34^. HMM profiling of MAEPL and MADA motifs across plant NLRs supports their divergence, with each motif enriched in their respective lineage. Nevertheless, MAEPL motifs from non-flowering plants functionally replaced the MADA motif of the angiosperm helper NLR NbNRC4 despite >450 million years of divergence. We hypothesize that MAEPL and MADA are derived from an ancestral motif, as phylogenetic analysis of CC_OG3_-NLRs and angiosperm CC-NLRs supports common ancestry. While each type of motif has diversified in overall sequence composition, conserved hydrophobic leucine residues are essential for non-flowering MAEPL activity similar to angiosperm MADA motifs^34^. The proper placement of hydrophobic residues within N-terminal helices therefore presents as a defining feature of CC-type executioner domains. This implies that sequence variation at the N-terminus can be accommodated provided that the appropriate distribution of hydrophobic residues is maintained. In agreement with this hypothesis, CC_RPW8_ executioner domains encode an N-terminal motif that is distinct from angiosperm CCs but retains similarly placed hydrophobic residues^19^. It is therefore conceivable that a range of N-terminal motifs have evolved within the diverse CC domains of plants. Precise biochemical mechanisms underpinning cell death activity remain to be clarified, though several lines of evidence point towards their involvement in the targeted disruption of cellular membranes and ion channel activity^15,17,34,49^. Further dissection of N-terminal motif evolution is therefore critical for revealing keystone molecular mechanisms of plant CC-NLR-mediated immunity.

We hypothesize that CC_OG3_-type NLRs carrying MAEPL function as singleton or helper NLRs in non-flowering lineages as is suggested for MADA-CC-NLRs^34^. By contrast, a prominent signature of sensor NLRs is the presence of integrated domains that bait pathogen effectors. In angiosperms, NLR-IDs form 5-10% of the NLRome_6_. Here, we identified a varying range of NLR-IDs across diverse NLR subtypes within ∼4% of the non-flowering plant NLRome. We failed to detect NLR-IDs in MAEPL-CC-NLRs, further supporting the idea that they function as singleton or helper NLRs. The integrated domains of non-flowering plants show similarity to angiosperm NLR-IDs and include protein domains known to be targeted by pathogen effectors like kinases, zinc fingers, thioredoxins, and transcription factors^6,50^. Moreover, non-flowering NLR-IDs were incorporated into broadly distributed (CC-NLRs & TIR-NLRs) as well as lineage-specific (Hyd-NLRs & Pkn-NLRs) immune receptors. At present, direct evidence for pathogen-induced NLR-mediated immunity is limited to angiosperms and remains to be discovered in non-flowering lineages. Whether MAEPL-CC-NLRs serve as helpers to activate immunity upon the perception of effectors by diverse sensor NLRs is an intriguing starting point for the future dissection of NLR-mediated immunity in non-flowering plants.

In *Marchantia*, MAEPL CC accumulation activated an immune-like response that included ion leakage and tissue browning that is often associated with oxidative and/or biotic stress in bryophytes^51–53^. Transcriptome analysis revealed a strong induction of pathogenesis-related and phenylpropanoid biosynthesis genes that are characteristic of induced defenses in *Marchantia*^40,54^. Further comparisons between *Marchantia* thalli and *Nicotiana* leaves provided our first look into shared CC responses in divergent plants. Here, the downregulation of cell homeostasis and chromatin-associated machinery was a common consequence of CC activation. Intriguingly, this included MiniChromosome Maintenance (MCM) complex genes involved in DNA replication. In metazoans, depletion of MCM abundance is associated with DNA replication failure leading to apoptosis^55,56^. In plants, MCM depletion causes ovule abortion and activation of the DNA damage response^57,58^. Consistent with these observations, direct activation of the CC-NLR RPM1 induces DNA damage in the model angiosperm *A. thaliana*^59^. Collectively, these observations suggest that the activation of NLR-mediated immunity perturbs DNA replication homeostasis leading to DNA damage and cell death. Whether this represents an ancestral component of the CC-NLR immune response remains to be clarified.

The broadly distributed TIR domain has a deep evolutionary history and contributes to immunity in plants, animals, fungi, oomycetes, bacteria, and archaea^5^. Indeed, functional interrogation of TIR domains has demonstrated conserved activity across angiosperms, as monocot TIR-only proteins induce HR cell death in the dicots *N. benthamiana* and *N. tabacum*^32,60^. Moreover, animal and bacterial TIRs display NADase activity, produce immune-related small molecules, and can activate HR cell death in plants (*Nicotiana*)^60,61^. Conversely, plant TIR domains exhibit conserved NADase activity in *Escherichia coli* similar to prokaryotic TIRs^60,62^. Our functional interrogation of TIR domains from non-flowering plants and streptophyte algae further confirms wide functional conservation. Intriguingly, this occurs even though seed-free plants lack EDS1, the central regulator of TIR activity in flowering plants^63^. A recent study demonstrated that EDS1 is not required for the activity of a monocot (maize) TNP-family TIR domain in *Nicotiana*^32^. Given that TNPs and TNLs originate in lineages lacking EDS1, it is likely that EDS1-independent immune responses arose early in the evolution of plants and algae. Further research exploiting non-flowering model systems enriched with TIRs (mosses) is likely to provide novel insights into the diversification of TIR-mediated immune mechanisms in plants.

Molecular genetic NLR research has historically focused on a limited group of angiosperm crops and model systems. Here, we took a comparative macroevolutionary approach to broaden our functional understanding of the NLR landscape in diverse plant lineages. We identified functionally conserved TIR and CC N-terminal domains spanning the spectrum of plant evolution. Our first look into CC functionality in the divergent model *M. polymorpha* revealed similarities with the angiosperm *N. benthamiana* despite over 450 million years of divergence between each system. Together, this suggests that NLRs are a core component of the plant immune system. While TIR and CC-type NLRs are likely to retain the functions of their angiosperm counterparts, non-flowering lineages harbor an untapped diversity of atypical NLRs with novel N-terminal executioner domains (hydrolases and kinases), C-terminal repeats, and integrated domains. This extended set of plant NLRs therefore provides exciting potential to discover novel mechanisms of plant disease resistance and to further dissect the fundamental aspects of immune receptor form and function. Indeed, conceptually similar studies exploring NLR-like proteins of bacteria are beginning to uncover such novelties^64^. Future studies aimed at exploiting the diverse receptor repertoires of plants and algae are therefore likely to advance our fundamental understanding of NLR-mediated immunity and may ultimately inform efforts to engineer novel disease resistance mechanisms in crops.

## ACKNOWLEDGMENTS

We thank Takayuki Kochi (Kyoto University, Japan) for providing pMpGWB168; Sebastian Schornack (Sainsbury Laboratory University of Cambridge, U.K.) and Tolga Bozkurt (Imperial College London, U.K.) for commenting on an earlier draft of the manuscript; our colleagues at the Joint Genome Institute for access to *Ceratodon, Ceratopteris, Physcomitrium*, and *Selaginella* genomic information (https://phytozome.jgi.doe.gov); Adeline Harant and Mauricio Contreras for technical support; Kristina Grenz, Kayla Robinson, Hyeonmin Jeong, Max Jordan, and all former members of the Carella group for additional technical and critical support. We thank the Department of Energy Joint Genome Institute, Jonathan Shaw (Duke University), and David Westin (ORNL) for pre-publication access to the proteomes of *Sphagnum fallax* and *S*.*magellanicum* that we used for NLR prediction. We also thank Michal Lorenc, Jiyuan Yan, and Peter Waterhouse (Queensland University of Technology, Australia) for pre-publication access to the *N. benthamiana* v3.5 draft genome (these sequence data were produced by the Nicotiana benthamiana Sequencing Consortium). This work was supported by the UKRI Biotechnology and Biological Sciences Research Council (BBSRC; grants BB/P012574, BBS/E/J/000PR9798, BBS/E/J/000PR9797, BBS/E/J/000PR9796, BBS/E/J/000PR9795) and the Gatsby Charitable Foundation.

## AUTHOR CONTRIBUTIONS

K.S.C, J.K., S.K., and P.C. designed research; K.S.C., and P.C. performed research; K.S.C., J.K., T.S., M.V, and P.C. analyzed data; K.S.C and P.C. prepared all figures and final datasets; P.C. wrote the paper with contributions from all authors.

## METHODS

### Plant Growth Details

*Marchantia polymorpha* (TAK1 background) were cultivated axenically from gemmae and grown under a long day photoperiod (16 hours of light; ∼80 μE light intensity) on one-half–strength MS (Murashige and Skoog) media (pH 6.7) with B5 vitamins at 20-22 °C. *Nicotiana benthamiana* were grown in soil under controlled conditions with a temperature of 22 °C and a long day photoperiod (16 hours of light; ∼160-200 μE light intensity).

### Confocal Microscopy

For experiments using *M. polymorpha*, confocal laser scanning microscopy was performed on a Leica TCS SP8 equipped with HyD detectors. A white light laser was used to visualize eYFP (excitation 515 nm) and mScarlet (excitation 570 nm). For experiments using *N. benthamiana*, confocal microscopy was performed on a Zeiss LSM880. Argon Ion (457 / 488 / 514 nm) and HeNe lasers (594nm) were used to visualize eYFP (excitation 515 nm) and RFP (excitation 594 nm). We collected images from at least three independent plants per experimental replicate. All experiments were performed at least three times with similar results.

### Trypan blue staining

Trypan blue staining was performed on liverworts using the protocol described in Redkar et al. (2022)^54^. In our conditions, the differential staining of stressed-vs-control liverworts was particularly difficult to discern since apical notches are easily stained and estradiol induction of CC activity led to tissue darkening and phenolic deposits in this area. Tissue clearing (chloral hydrate) of unstained liverworts verified this pattern. Trypan blue staining of heat-killed liverworts and *N. benthamiana* leaves undergoing CC-eYFP-induced HR confirmed the viability of l staining solutions. For *N. benthamiana*, trypan blue staining was performed as described in Ma et al. (2012)^42^.

### Transient Agrobacterium-mediated expression and cell death assays

Transient expression of all constructs in *Nicotiana* were performed by agroinfiltration according to methods described in Adachi et al. (2019)^34^. Briefly, four weeks old *Nicotiana* plants were infiltrated with Agrobacterium tumefaciens GV3101 carrying binary expression plasmids. *A. tumefaciens* suspensions were prepared in fresh infiltration buffer (10 mM MES-KOH,10 mM MgCl2, and 150 mM acetosyringone, pH5.6) and adjusted to OD_600_ = 0.25 that were then mixed in a 1:1 ratio with an *A. tumefaciens* carrying the p19 silencing suppressor. HR cell death phenotypes were scored on an HR index ranging from 0 (no visible symptoms) to 7 (fully confluent cell death).

### Estradiol-induction and ion leakage assays

Estradiol-inducible gene expression in *Marchantia* was achieved by vacuum infiltration generated within the cavity of a needless 50 mL syringe. A 20 mM stock of estradiol (in 100% DMSO) was used to prepare a 50 or 20 μM working concentration (in water). For mock-treatment controls, we used a comparable 0.25% or 0.1% DMSO solution (in water). To minimize damage caused by tissue handling, all liverworts used for infiltrations were grown on nylon mesh (Normesh, UK), which allowed easy transfer of liverwort thalli to the syringe for treatment. Once infiltrated, thalli were subsequently transferred to clean petri dishes containing at least 2 layers of pre-wetted filter paper. Plates were sealed with micropore tape and returned to the appropriate growing condition. Where indicated, estradiol treatment was also performed by culturing liverwort gemmae directly into solid media supplemented with estradiol (20 μM) or DMSO (0.1%). Ion leakage assays were performed essentially as described in Hatsugai and Katagiri (2018)^43^. Measurements were performed after 4, 24 and 48 hours post-harvest with a compact conductivity meter (LAQUAtwin-EC-33, Horiba Scientific). Five leaf discs (4mm diameter) were harvested from different thallus and immerse inside 2ml ddH2O as one biological replicate. Each measurement contains three biological replicates and all experiments were performed at least three times.

### RNA Isolation, cDNA Synthesis, and qRT-PCR Analysis

Total RNA was extracted from flash-frozen *M. polymorpha* (TAK1) plants that were collected 24 hours post mock (0.1% DMSO in water) or estradiol (20 μM) treatment using the Spectrum Plant RNA Extraction Kit (Protocol A) with on-column DNAse treatment following the manufacturer’s instructions. cDNA was synthesized using 2 μg of total RNA with SuperScript II reverse transcriptase (Invitrogen) following the manufacturer’s instructions. qRT-PCR reactions were performed in a total volume of 10 μL using 2.5 μL of 10x-diluted (in nuclease-free water) cDNA and Roche SYBR mix with the primers listed in Data S6. The qRT-PCR reaction protocol consisted of an initial denaturation at 95 °C for 5 minutes followed by 40 cycles of 95 °C for 10 seconds, 60 °C for 15 seconds, and 72 °C 15 seconds on a Roche LightCycler 480 II according to manufacturer’s instructions. All qRT primers were designed using Primer3^65,66^ or were previously published as listed in Data S6. Specificity was validated by analyzing melt curves after each run. Three independent sample replicates as well as three technical replicated per sample were performed at any given time point/treatment. Calculations of expression levels normalized to internal controls and statistical analyses (ANOVA, Tukey’s HSD) were performed using R software and all graphs were generated in GraphPad Prism (v9.3.1).

### Cloning and Marchantia transformation

NbNRC4 chimera constructs were amplified by PCR with primers containing BsaI cloning sites, the appropriate N-terminal motif, and overlapping NbNRC4^D478V^ sequences using a NbNRC4^D478V^-6HA construct^34^ as a template (Data S6). To generate NLR N-terminal domain (CC and TIR) eYFP fusion constructs, we synthesized codon optimized domains flanked by BsaI restriction sites (Azenta). Synthesized gene fragments, or the β-glucuronidase (GUS) control (Addgene #50332), were assembled with pICH85281 [mannopine synthase promoter+W (MasWpro), Addgene no.50272], pICSL50005 (YFP, TSL SynBio), pICSL60008 [Arabidopsis heat shock protein terminator (HSPter), TSL SynBio] and the binary vector pICH47742 (Addgene no. 48001) in a Golden Gate compatible system. To generate Marchantia expression vectors, the MpCNL1 (Mp3g01950,1-266aa) and MpCNL1ΔN (Mp3g01950,31-266aa) were cloned by PCR (Q5 High Fidelity Polymerase, NEB) with attL-containing primers using codon optimized gene fragments as a template. mCitrine-HA and myr-mScarlet were amplified from template plasmids containing the respective fluorophore; mCitrine/pMpGWB105 (Addgene # 68559) and mScarlet:CETN2^67^. To generate fluorophore fusions, we used a multi-step PCR approach with overlapping primers to generate an HA tag on the C-terminal end of mCitrine, or a myristolation sequence at the N-terminus of mScarlet. Amplicons were flanked by attB sites to enable recombination into pDONR221 using BP Clonase II (Invitrogen) following manufacturer instructions. MpCNL1^CC^-eYFP, MpCNL1^CCΔN^-eYFP and mCitrine-HA inducible expression constructs were generated by LR recombination reactions into pMpGWB168 (XVE::GW)^39^. The plasma-membrane marker construct MpEF1a:myr-mScarlet was generated by LR recombination into pMpGWB303 (Addgene no. 68631)^68^. All resulting constructs were transformed into *Agrobacterium tumefaciens* GV3101 (pMP90) by electroporation. *M. polymorpha* transformation was performed using the *Agrobacterium*-mediated thallus regeneration method^69^ in the TAK1 background. Transformants were selected on solid one-half strength MS-B5 media supplemented with cefotaxime (125 μg/mL) and hygromycin B (15 - 25 μg/mL) or chlorsulfuron (0.5 - 1 μM). Stable transgenic plants were obtained by propagating gemmae from T1 thalli. All experiments were performed in the G2 (second asexual/gemmae generation) or subsequent generations.

### Protein immunoblotting

Protein samples were prepared from four tissue discs (8 mm diameter) sampled from *N. benthamiana* leaves at 1 day or 2 days after agroinfiltration and were homogenized in 100 μL of 2X SDS loading buffer (0.1 M Tris-HCl, 0.2 M DTT, 4.0% [w/v] SDS, 3mM Bromophenol blue, 2 M Glycerol). Protein samples from *Marchantia* were prepared from five tissue discs (4mm diameter) with 50 μL of 2X SDS loading buffer. Immunoblotting was performed with HA-probe (F-7) HRP (sc-7392 HRP, Santa Cruz Biotech, 1:5000 dilution) or primary anti-GFP antibody (11814460001, Roche, 1:2500 dilution) combined with secondary HRP-linked Anti-mouse IgG (NXA931-1ML, Amersham,1:5000 dilution). Total protein loading was visualized by staining nitrocellulose membranes with Ponceau S solution (Sigma-Aldrich, P7170).

### Library preparation and sequencing

mRNA from *M. polymorpha* transgenics 24 hours post estradiol-treatment or *N. benthamiana* leaves 24 hours post agroinfiltration were purified from DNAse-treated total RNA (prepared as described above) using Poly(A) selection and then fragmented (at least 3 independently treated plants collected per sample replicate). cDNA library preparation was performed with the TruSeq® RNA Sample Preparation Kit (Illumina, US) according to manufacturer’s instructions. Sequencing of each sample (in triplicate) was performed on the Illumina NovaSeq in 150 paired end mode. De-multiplexed samples were used for subsequent analyses. All raw fastq data are accessible at http://www.ncbi.nlm.nih.gov/sra/ under the accession number PRJNA881591.

### Expression analysis

We first analyzed raw sequencing reads with FastQC for quality control (https://www.bioinformatics.babraham.ac.uk/projects/fastqc/). Reads were then aligned back to the appropriate genome (*Marchantia polymorpha* v5.1 - https://marchantia.info/download/tak1v5.1/; *Nicotiana benthamiana* draft genome v3.5 - https://www.nbenth.com/) using HiSAT2^70^. We used featureCounts^71^ to obtain raw counts using only uniquely mapped and properly paired reads. Differentially expressed genes were identified with DESeq2^72^ following pair-wise comparisons between controls (mCitrine-HA control for *Marchantia* experiments; GUS-YFP control for *N. benthamiana* experiments) and the indicated treatment conditions. We focused only on differentially expressed genes with a strict cut-off (absolute LFC [log2 fold change] ≥ 2 and adjusted p-value ≤ 10^−3^) when performing hierarchical clustering of samples. Heatmaps were generated with R pheatmap using variance-stabilised counts median-centered by gene. Functional enrichment analyses were performed using the MBEX online tool (https://marchantia.info/mbex/)^73^ with a significance cutoff of FDR (False Discovery Rate) ≤ 0.05.

### NLR prediction, phylogenetics, orthology, and protein sequence analyses

We used NLRtracker to identify and annotate NLRs in the genome annotations listed in Data S1 (sheet 1). For phylogenetic analysis, we first generated amino acid sequence alignments using only the NB-ARC domains of predicted NLRs using MAFFT^74^. After trimming the alignment with trimAl^75^, we used IQ-TREE (v2.0.3)^76^ to perform maximum likelihood phylogenetic analysis with bootstrapping (1000 Ufboot + SH-aLRT). The ModelFinder option identified ‘JTT+F+G4’ as the optimal substitution model for our dataset. All subsequent tree rendering was performed using iTOL, and a public tree with extended annotation options can be found online (https://itol.embl.de/shared/philcarella).

We used OrthoFinder (OrthoFinder-2.5.4)^45^ to reconstruct orthologous protein groups (orthogroups) shared between organisms. For orthology analysis of NLR N-terminal domains we considered any protein sequence upstream of the NB-ARC domain as a putative N-terminal domain and performed OrthoFinder on de-duplicated sequences. For comparative transcriptomics, we performed OrthoFinder on the proteomes (primary protein isoforms only) of *M. polymorpha* and *N. benthamiana*. Visualization of differentially expressed orthologs was performed using GraphPad Prism9 software. Single-copy orthologs were compared 1-to-1, whereas multi-copy orthologs were compared 1-to-many or many-to-1 (i.e. the same *Marchantia* value may be compared to multiple *Nicotiana* values and *vice versa*).

To identify conserved amino acid motifs in OG3-type CCs, we subjected an amino acid sequence alignment of CC_OG3_ domain sequences to MEME analysis^33^ with the parameters ‘zero or one occurrence per sequence, top ten motifs’. MEME detected the N-terminal MAEPL motif in several CC_OG3_ domains, therefore we curated a more specific alignment containing these N-terminal sequences to build a hidden Markov model (HMM) profile of the non-flowering ‘MAEPL’ motif using the ‘hmmbuild’ function in HMMER (version 3.3.2)^35^. Upon calibration with ‘hmmcalibrate’, we tested the MAEPL.hmm profile in the proteomes of major plant lineage representatives listed in Data S1. We further compared the MAEPL.hmm with the angiosperm MADA.hmm profile^34^ in two representative angiosperm proteomes; *A. thaliana* (TAIR10, https://www.arabidopsis.org/) and *S. lycopersicum* (ITAG4.1, https://solgenomics.net/organism/Solanum_lycopersicum/genome) and in the angiosperm NLR atlas (https://biobigdata.nju.edu.cn/ANNA/)^31^. All motif scans were performed using the following template line of code ‘hmmsearch --max -o output.txt Motif.hmm Proteome.fasta’.Supporting files and raw data related to NLR prediction, NB-ARC phylogenetic analysis, N-terminal domain orthology, N-terminal domain structural prediction, and HMM profiling are deposited at Zenodo doi: 10.5281/zenodo.7092643.

### Statistics

Statistical details of experiments can be found in the corresponding figure legends. Here, the identity of the statistical tests used, the exact value of n (i.e. number of independently infected liverworts) and dispersion and precision measures are given (error bars represent mean +/-standard deviation, p-value cutoffs, etc.). All statistical analyses for transcriptomic and proteomic analyses are described in the methods details above. Statistical analysis of qRT-PCR expression data are described in figure legends and were performed using R. Student’s t-tests were performed in Microsoft Excel or GraphPad Prism.

## SUPPLEMENTARY INFORMATION

**Figure S1. Predicted structures of NLR N-terminal executioner domains**

**Figure S2. MAEPL motif occurrence across major plant lineages**

**Figure S3. Subcellular localization of MAEPL-CC_OG3_ in *Nicotiana* and *Marchantia***

**Figure S4. Characterization of the MAEPL-CC_OG3_ response in *Marchantia***

**Figure S5. Transcriptomic analysis of CC_OG3_ activity in *Marchantia* and *Nicotiana***

**Data S1. NLR prediction and orthology analysis**

Data **related to Figure 1 and Figure 2**. Includes a full list of queried genomes (sheet 1), characterization of NLRs identified through NLR tracker (sheets 2-4), and N-terminal domain orthology analysis (sheet 5).

**Data S2. Cell death phenotypes of NLR N-terminal executioner domains in *Nicotiana***

Data **related to Figure 3 and Figure 4**. Macroscopic HR cell death phenotypes (A) indicative of the HR cell death index, (B) TIR-eYFP fusions in *N. benthamiana*, (C) TIR-eYFP fusions in *N. tabacum*, (D) CC-eYFP and CC_RPW8_-eYFP fusions in *N. benthamiana*, € MAEPL CC N-terminal truncation-eYFP fusions in *N. benthamiana*, (F) MAEPL L-to-E variant-eYFP fusions in *N. benthamiana*, and (G) MADA/MAEPL NbNRC4-6HA chimeras in *N. benthamiana*.

**Data S3. Protein immunoblotting**

Data **related to Figure 3, Figure 4, Figure 5, Figure S3 and Figure S4**. Protein immunoblots of (A) TIR/CC-eYFP fusions expressed in *N. benthamiana*, (B) MADA/MAEPL NbNRC4-6HA fusions in *N. benthamiana*, (C) MAEPL truncation/variant-eYFP fusions in *N. benthamiana*, and (D) *Marchantia* XVE transgenic lines.

**Data S4. MAEPL motif discovery**

Informatics data related to **Figure 4 and Figure S2**. Support for MAEPL discovery through (A) MEME motif identification, (B) MEME motif location, (C) MAEPL motif amino acid sequence alignment in OG3-type CCs, and (D) MAEPL/MADA comparison through amino acid sequence alignment.

**Data S5. RNA-sequencing analysis of CC_OG3_ activity in *Marchantia* and *Nicotiana*** Transcriptomics data **related to Figure 6 and Figure S6**. For sheets (1-2;4-6): significantly differentially expressed genes during CC induction (adjusted p-value < 10^−3^, log fold change (|LFC| ⩾2), variance-stabilized row-centered counts are shown.

**Data S6. Oligonucleotide primers and synthetic gene fragments used in this study**

