## Supplementary Datasets for "The N-terminal executioner domains of NLR immune receptors are functionally conserved across major plant lineages": DataS2.pdf

**Data S2**

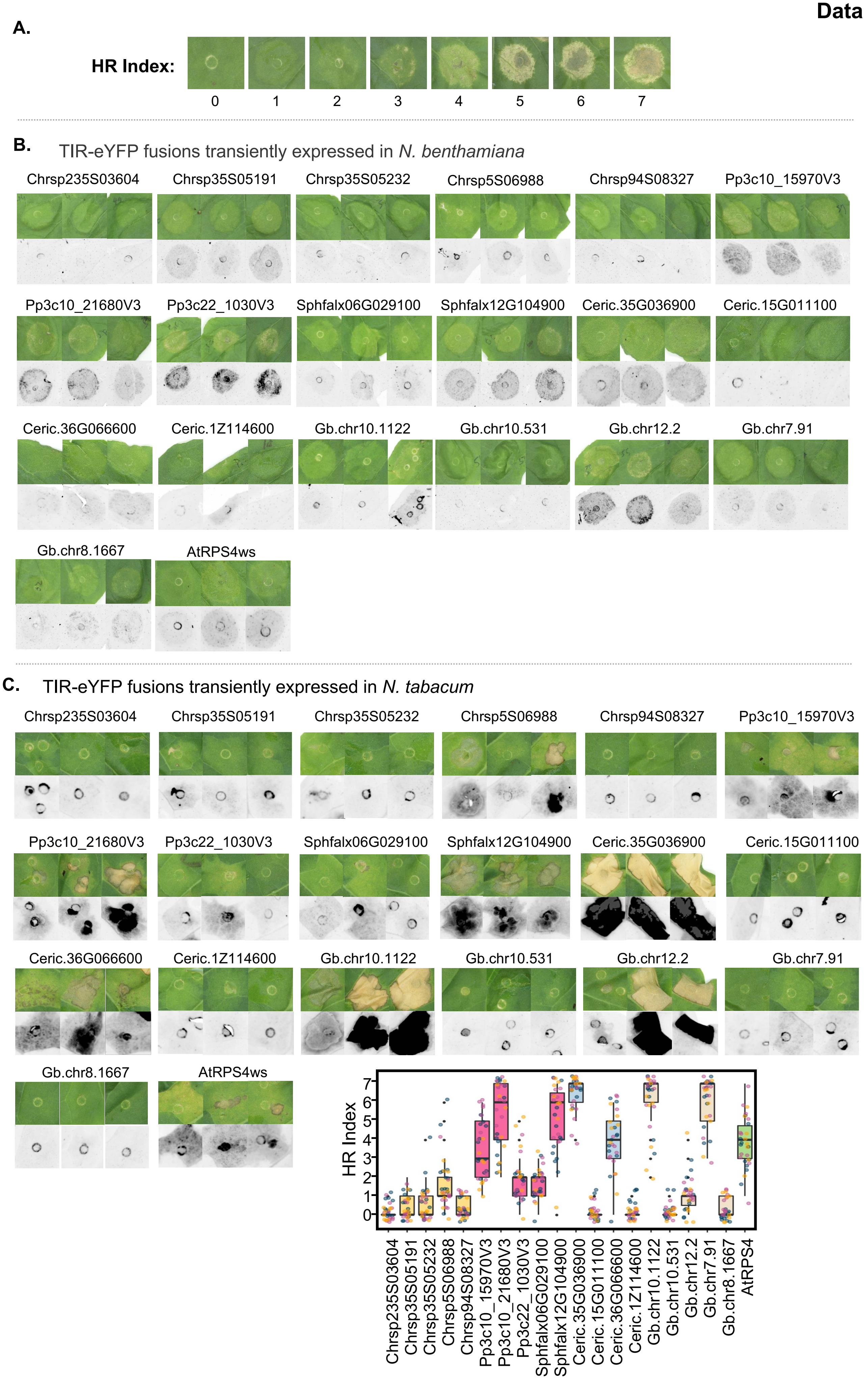
A.
HR Index:
0
1
2
3
4
5
6
7
B.
TIR-eYFP fusions transiently expressed in N. benthamiana
Chrsp235S03604
Chrsp35S05191
Chrsp35S05232
Chrsp5S06988
Chrsp94S08327
Pp3c10_15970V3
Pp3c10_21680V3
Pp3c22_1030V3
Sphfalx06G029100
Sphfalx12G104900
Ceric.35G036900
Ceric.15G011100
Ceric.36G066600
Ceric.1Z114600
Gb.chr10.1122
Gb.chr10.531
Gb.chr12.2
Gb.chr7.91
Gb.chr8.1667
AtRPS4ws
C.
TIR-eYFP fusions transiently expressed in N. tabacum
Chrsp235S03604
Chrsp35S05191
Chrsp35S05232
Chrsp5S06988
Chrsp94S08327
Pp3c10_15970V3
Pp3c10_21680V3
Pp3c22_1030V3
Sphfalx06G029100
Sphfalx12G104900
Ceric.35G036900
Ceric.15G011100
Ceric.36G066600
Ceric.1Z114600
Gb.chr10.1122
Gb.chr10.531
Gb.chr12.2
Gb.chr7.91
Gb.chr8.1667
AtRPS4ws
HR Index
0
1
2
3
4
5
6
7
Chrsp235S03604
Chrsp35S05191
Chrsp35S05232
Chrsp5S06988
Chrsp94S08327
Pp3c10_15970V3
Pp3c10_21680V3
Pp3c22_1030V3
Sphfalx06G029100
Sphfalx12G104900
Ceric.35G036900
Ceric.15G011100
Ceric.36G066600
Ceric.1Z114600
Gb.chr10.1122
Gb.chr10.531
Gb.chr12.2
Gb.chr7.91
Gb.chr8.1667
AtRPS4

**D.** CC-eYFP & $CC_{RPW8}$-eYFP fusions transiently expressed in *N. benthamiana*

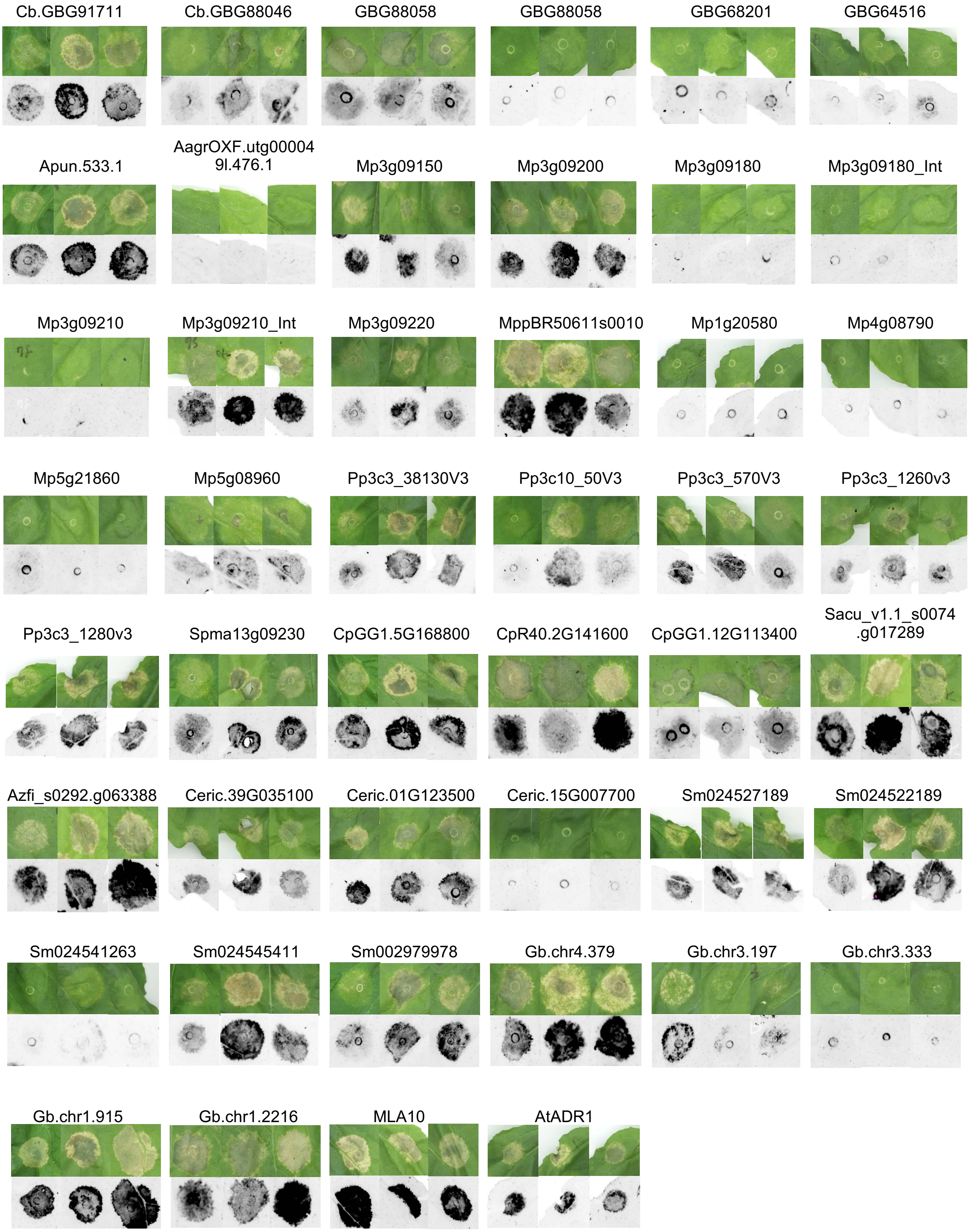

**E.** MAEPL-CC domains and N-terminal truncations fused with eYFP

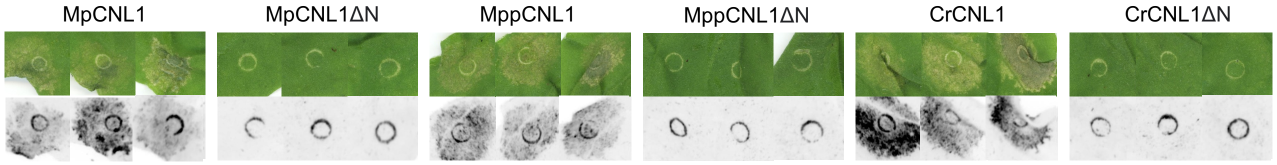


**F.** MAEPL-CC leucine variants (2E/3E) fused with eYFP

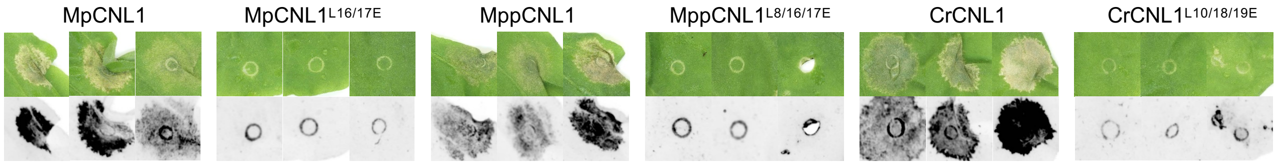


**G.** MADA/MAEPL chimera fused with autoactivated NbNRC4$^{D478V}$-6HA

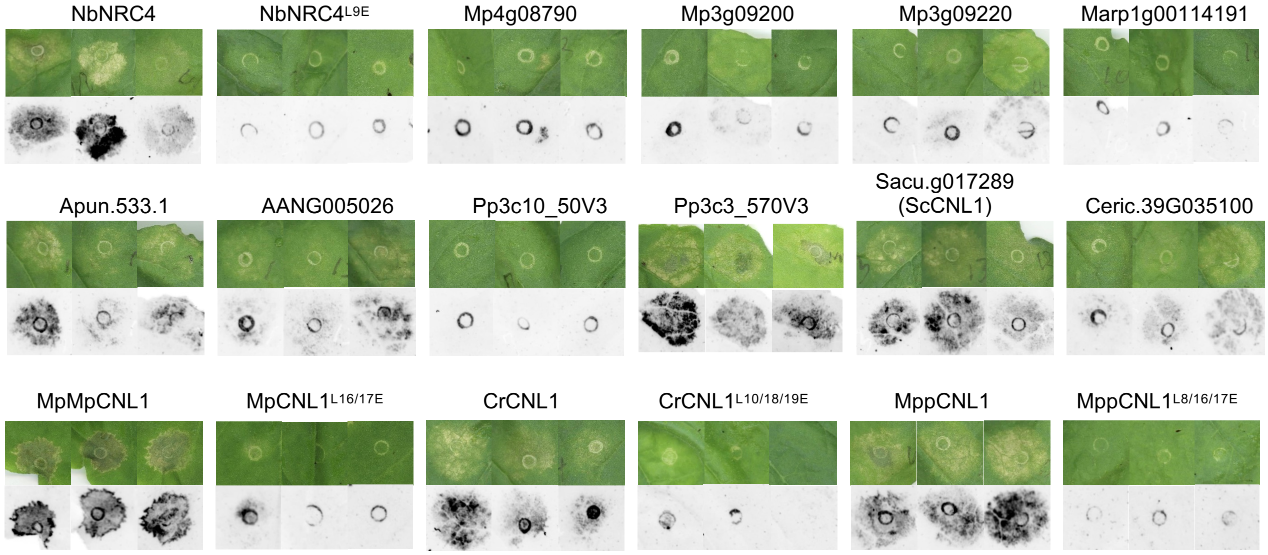


**Data S2. Cell death phenotypes of NLR N-terminal domains in *Nicotiana***

**(A)** Representative macroscopic cell death phenotypes illustrating the HR index (0-7) in *N. benthamiana.*

**(B)** Macroscopic phenotypes of TIR-eYFP fusion constructs at 5 days post infiltration (dpi) in *N. benthamiana.*

**(C)** Macroscopic phenotypes of TIR-eYFP fusion constructs at 5 dpi in *N. tabacum.* The HR index plot shows the distribution of cell death phenotypes across all three experimental replicates and was performed on the same scale as experiments utilizing *N. benthamiana* (Figure 3).

**(D)** Macroscopic phenotypes of CC-eYFP fusion constructs at 5dpi in *N. benthamiana.*

**(E)** Macroscopic phenotypes of wildtype and N-terminally truncated (ΔN) OG3 CC-eYFP fusion constructs at 5dpi in *N. benthamiana.* Gene symbols correspond to Mp3g09150 (MpCNL1), MppBR5_0611s0010 (MppCNL1), and Ceric.01G123500 (CrCNL1).

**(F)** Macroscopic phenotypes of wildtype and MAEPL variants (leucine to glutamic acid) of OG3 CC-eYFP fusion constructs at 5dpi in *N. benthamiana.*

**(G)** Macroscopic phenotypes of MAEPL chimera constructs at 5dpi in *N. benthamiana.* An autoactivated version of the MADA-type helper NLR NbNRC4$^{D478V}$-6HA was used as a scaffold to interrogate MAEPL function. Names indicate the source of the N-terminal motif used for each chimera. The MAEPL variants used here are described in (F).

All images (cropped and combined from independent leaves) are representative of three independent experimental replicates, with at least 9 infiltration sites per construct for each replicate. HR phenotypes are shown in bright-field and in UV. For experiments using *N. benthamiana,* the P19 suppressor was co-infiltrated to enhance transient expression. P19 was not utilized in experiments using *N. tabacum* since it activates cell death in this species.
