## Supplementary Datasets for "The N-terminal executioner domains of NLR immune receptors are functionally conserved across major plant lineages": DataS3.pdf

**Data S3**

**A.**

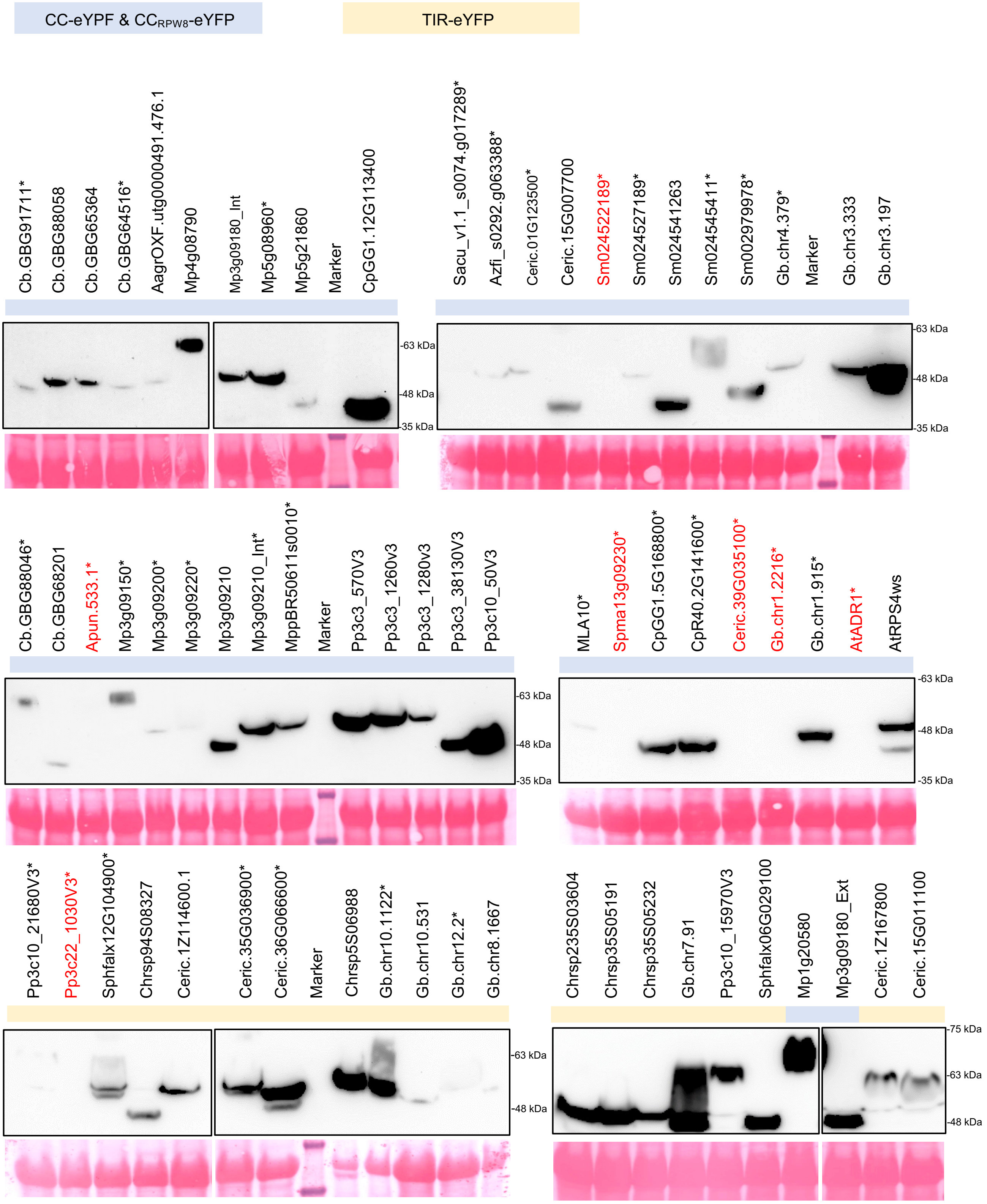


**Data S3. Protein immunoblotting**

(**A**) Anti-GFP immunoblots of NLR N-terminal domains (CC, $CC_{RPW8}$, and TIR) fused to eYFP. Total proteins were prepared from *N. benthamiana* leaves at 1 day after agroinfiltration for cell death inducing constructs or 2 days after agroinfiltration for the remaining constructs. Seven constructs activating strong cell death phenotypes (AtADR1, Ceric.39G035100, Sm024522189, Apun…533.1, Gb.chr1.2216, Spma13g09230, Pp3c22_1030V3) failed detection (names in red). Asterisks (*) indicate cell-death inducing constructs. Ponceau staining was used to visualize total protein loading.

### Data S3

**B.**

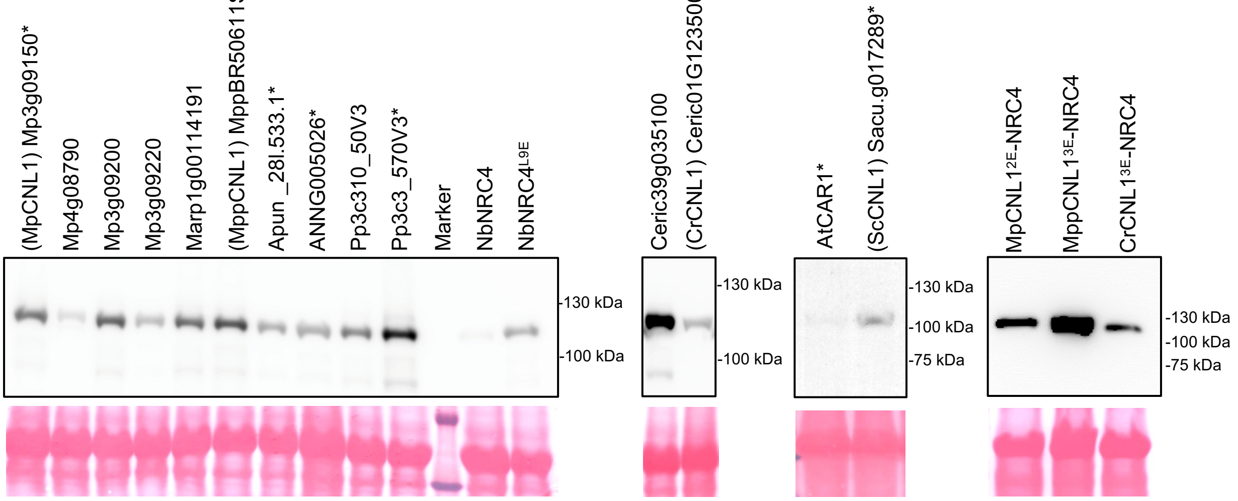


**C.**

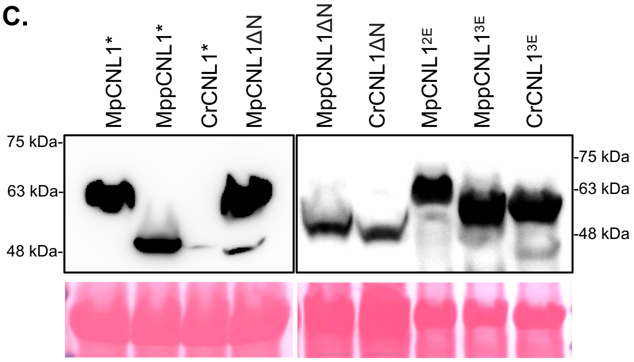


**D.**

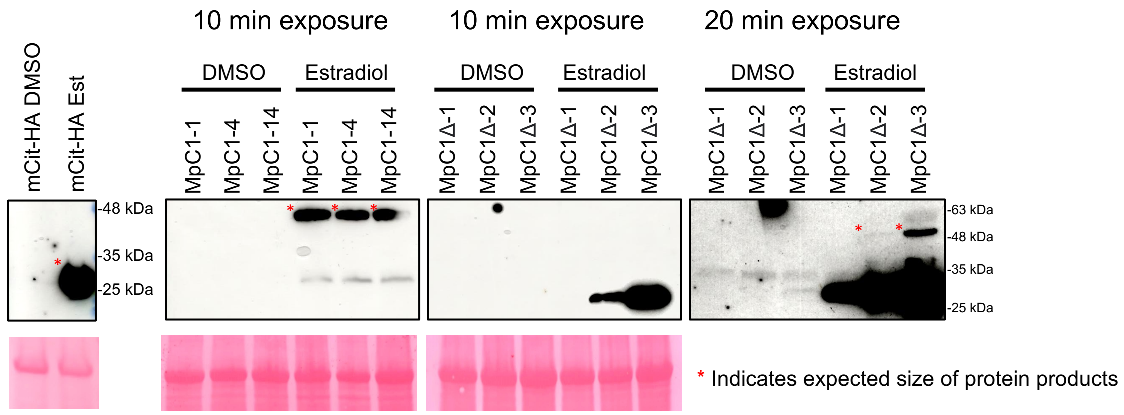


**Data S3 continued**

(**B**) Anti-HA immunoblots of NbNRC4$^{D478V}$-6HA variants/chimeras transiently expressed in *N. benthamiana* approximately 24-48 hours post agroinfiltration. Asterisks (*) indicate cell-death inducing constructs. Ponceau staining was used to visualize total protein loading.

(**C**) Anti-GFP immunoblots of N-terminally truncated and L-to-E mutated versions of MpCNL1, MppCNL1 and CrCNL1 domains fused with YFP transiently expressed in *N. benthamiana* approximately 24-48 hours post agroinfiltration. Asterisks (*) indicate cell-death inducing constructs. Ponceau staining was used to visualize total protein loading.

(**D**) Anti-GFP immunoblots of *Marchantia polymorpha* transgenic lines 24 hours post induction with either DMSO or Estradiol. Lines correspond to XVE:MpCNL1$^{CC}$-eYFP (MpC1), XVE:MpCNL1$^{CC\Delta N}$-eYFP (MpC1Δ), and XVE:mCitrine-HA (mCit-HA) in the TAK1 background. Ponceau staining was used to visualize total protein loading. For MpC1Δ lines, we provide exposures at 10 and 20 minutes to visualize expected fusion products (the single Ponceau loading control corresponds to both exposures).
