## Supplementary Datasets for "The N-terminal executioner domains of NLR immune receptors are functionally conserved across major plant lineages": DataS4.pdf

**A.**

| Motif | LOGO | E-value | Sites | Width |
|---|---|---|---|---|
| 1 | 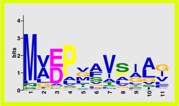 | 1.0e-161 | 79 | 11 |
| 2 | 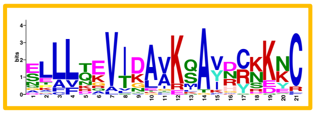 | 2.7e-280 | 38 | 21 |
| 3 | 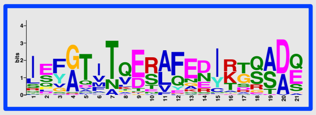 | 9.9e-295 | 44 | 21 |
| 4 | 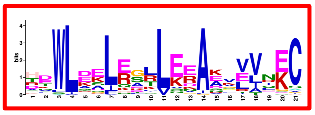 | 1.4e-542 | 75 | 21 |
| 5 | 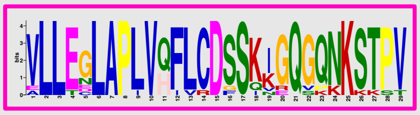 | 1.6e-214 | 13 | 29 |
| 6 | 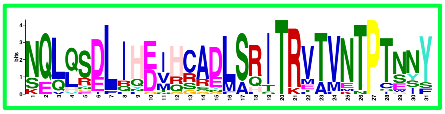 | 2.0e-295 | 19 | 31 |
| 7 | 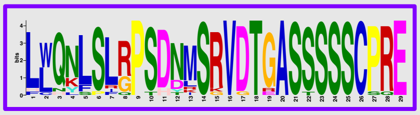 | 3.7e-313 | 17 | 29 |
| 8 | 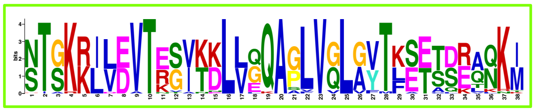 | 1.7e-615 | 29 | 38 |
| 9 | 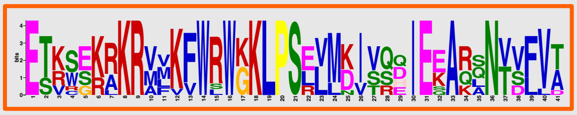 | 2.5e-179 | 10 | 41 |
| 10 | 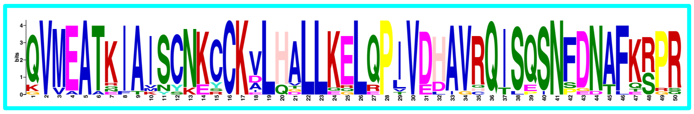 | 7.2e-644 | 18 | 50 |

**Data S4. MAEPL motif discovery**
(**A**) MEME analysis (MEMEsuite) using 80 OG3-type CC domains encoded within non-flowering land plants. The top ten amino acid motifs with “zero or one occurrence per sequence” are depicted. Width denotes size in amino acid residues.

**B.**

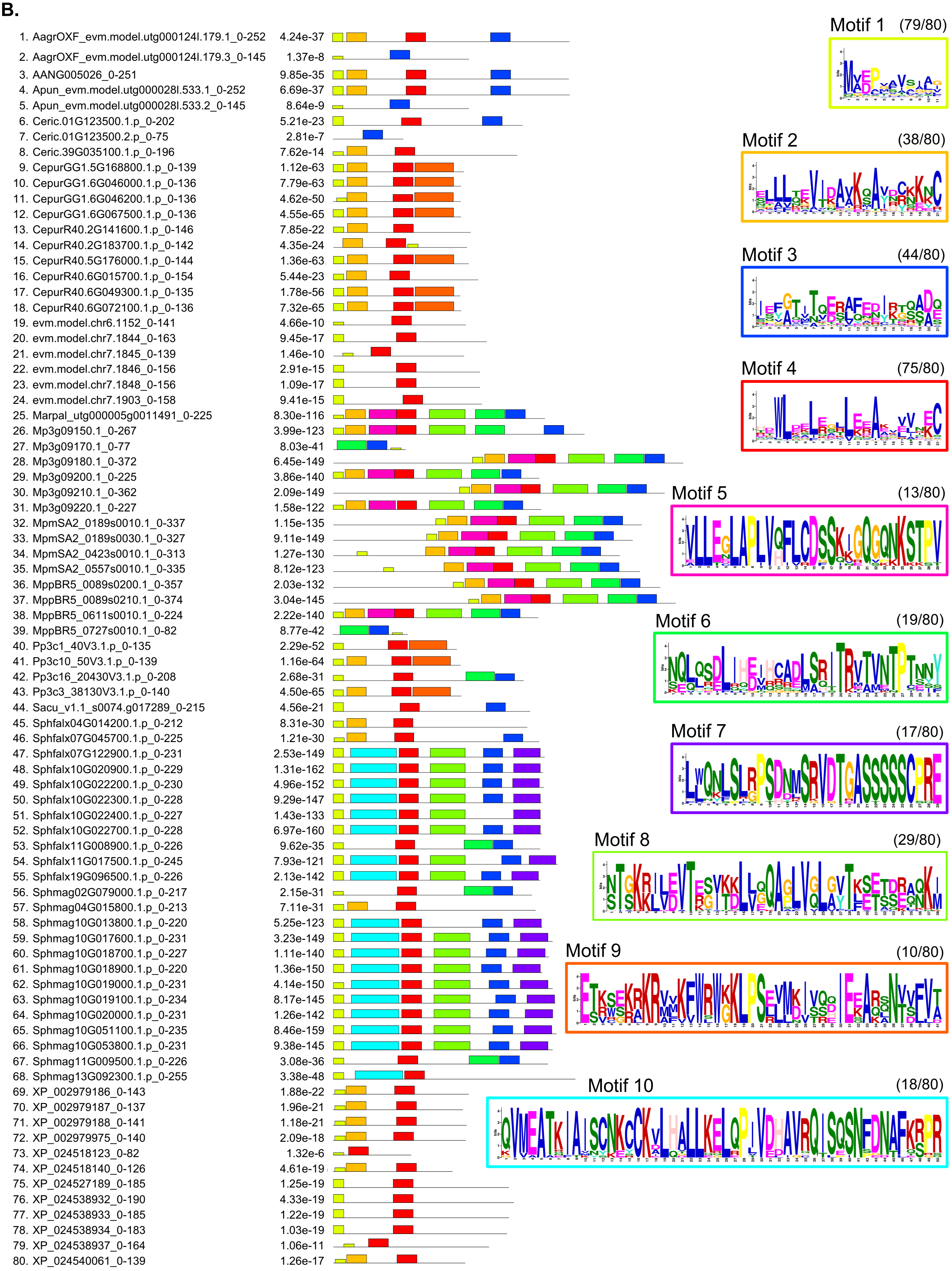


**Data S4. MAEPL motif discovery (continued)**

(**B**) Visual representations of motif location across 80 OG3 CC domains. The prevalence of each motif (x/80) is depicted next to each amino acid sequence LOGO.

**C.**

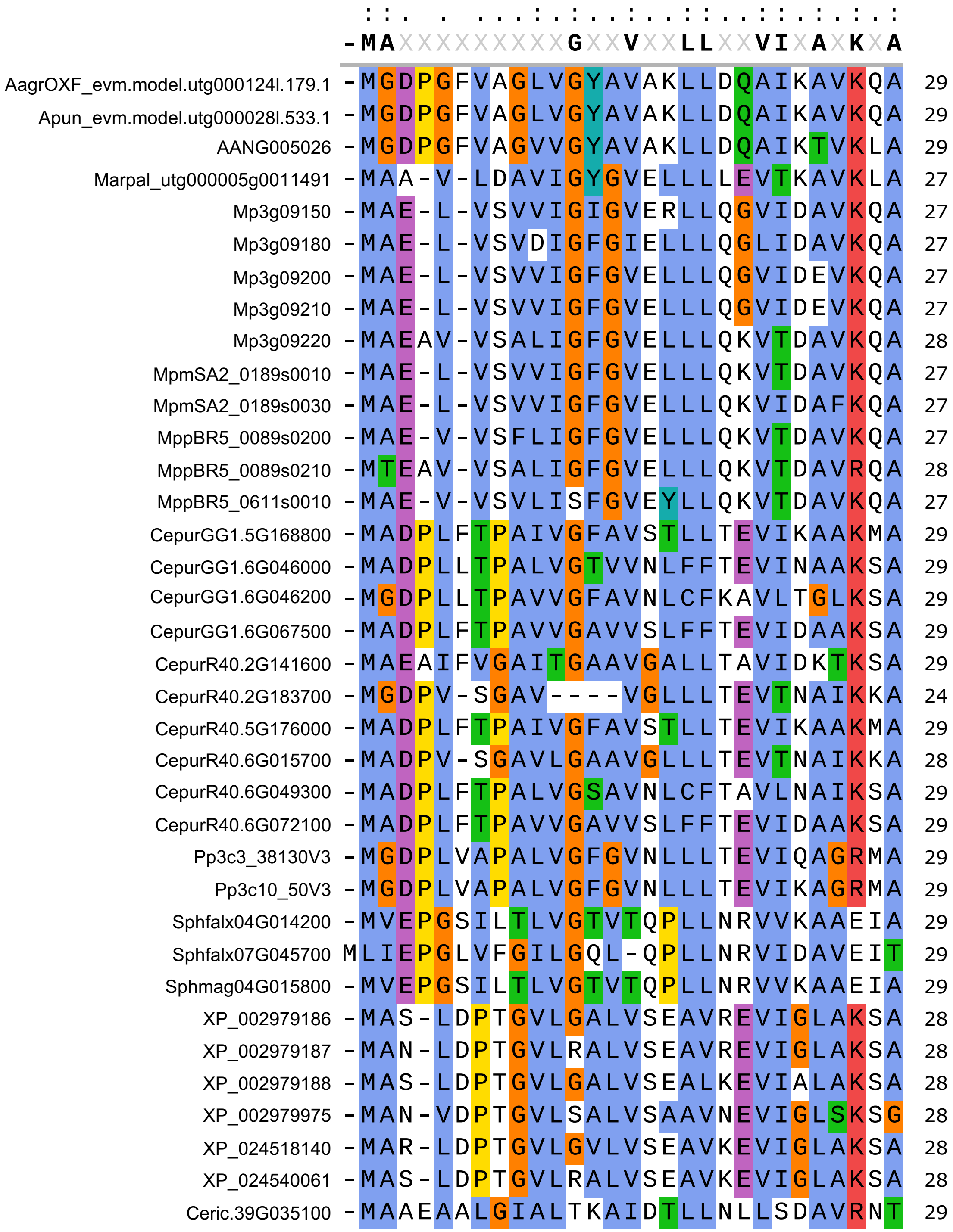


**Data S4. MAEPL motif discovery (continued)**

(**C**) Amino acid sequence alignment of the N-terminal MAEPL motif present in a subset of OG3-type CC domains of non-flowering plants. Alignments were performed using MAFFT in the SnapGene tool (v6.0.2). The consensus sequence shown represents residues present in >60% of all sequences. Coloring is based on amino acid residue properties + conservation (ClustalX).

**D.**

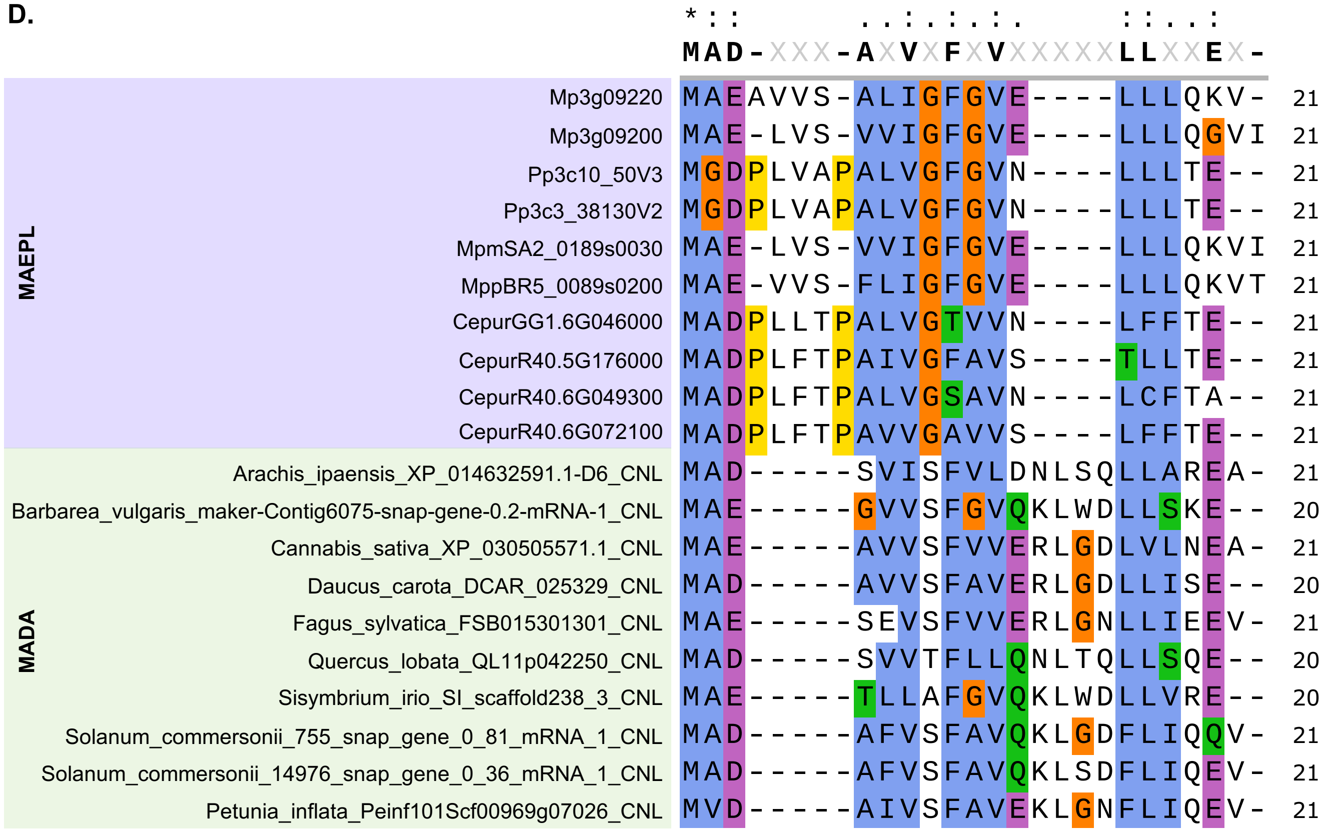


**Data S4. MAEPL motif discovery (continued)**
(**D**) Amino acid sequence alignment of 10 high-scoring MAEPL motifs (non-flowering OG3-type CC domains) and 10 high-scoring MADA motifs (queried against CC domains in the angiosperm NLR atlas). Alignments were performed using MAFFT in the SnapGene tool (v6.0.2). The consensus sequence shown represents residues present in >50% of all sequences. Coloring is based on amino acid residue properties + conservation (ClustalX).
