## Supplementary Datasets for "The N-terminal executioner domains of NLR immune receptors are functionally conserved across major plant lineages": FigureS1.pdf

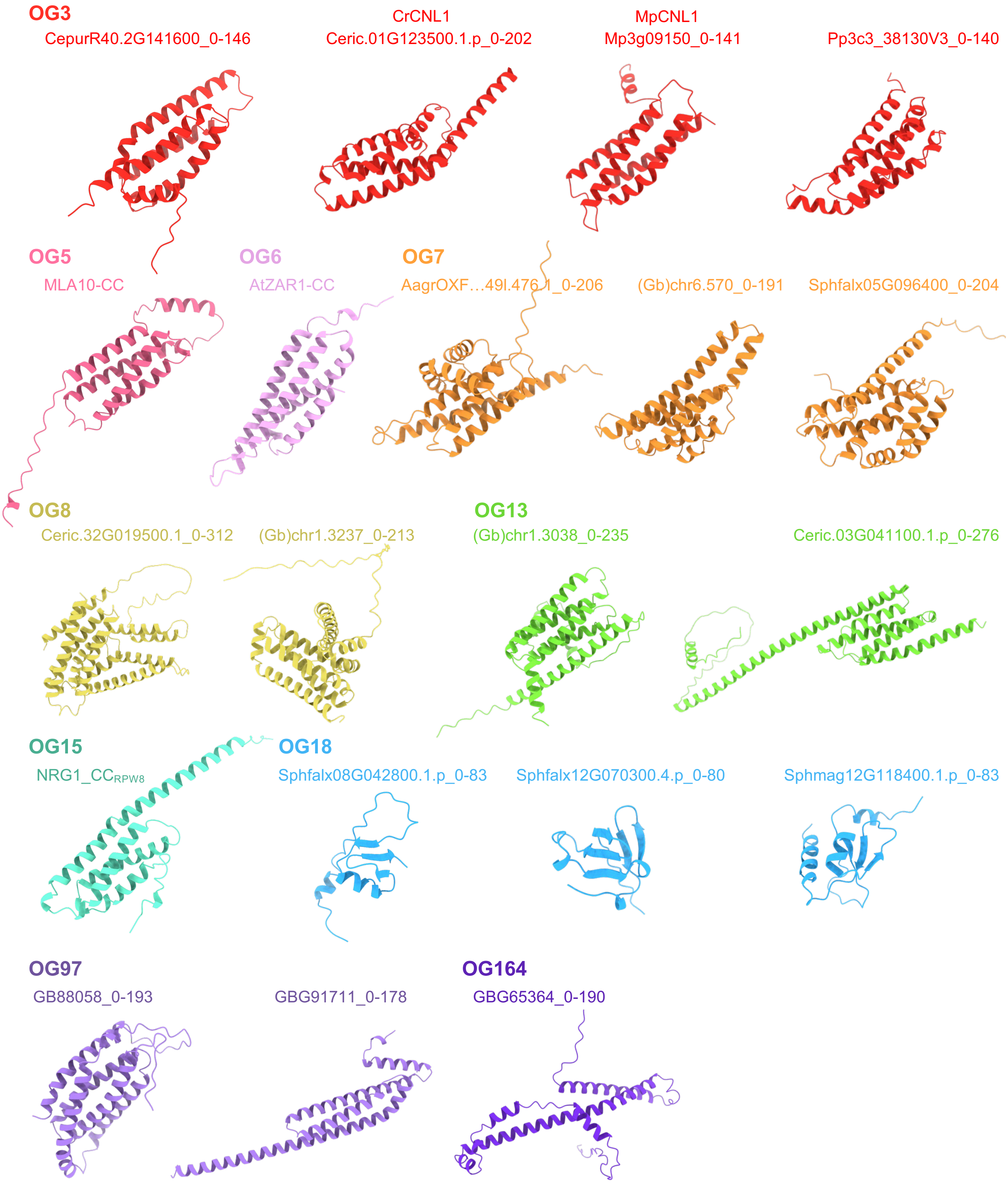


**Figure S1. Predicted structures of NLR N-terminal executioner domains**
AlphaFold2-based structural predictions of representative NLR N-terminal domains belonging to key orthogroups (OGs). Predictions were performed directly in ChimeraX-1.3 using the AlphaFold2 prediction (Collab-Fold) option. Each orthogroup is given a different color.
