## Supplementary Datasets for "The N-terminal executioner domains of NLR immune receptors are functionally conserved across major plant lineages": FigureS2.pdf

A.

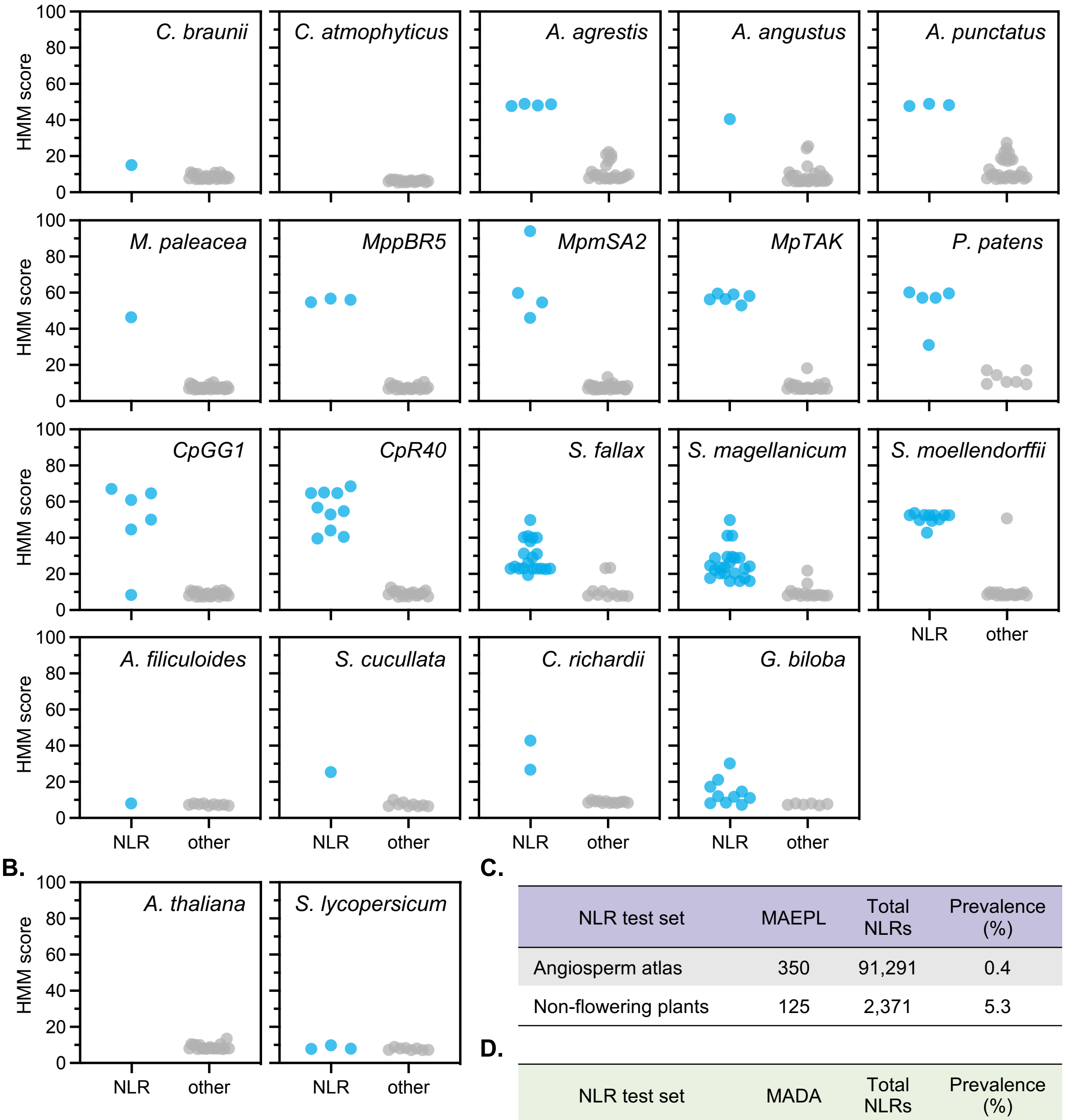


C.

| NLR test set | MAEPL | Total NLRs | Prevalence (%) |
|---|---|---|---|
| Angiosperm atlas | 350 | 91,291 | 0.4 |
| Non-flowering plants | 125 | 2,371 | 5.3 |

D.

| NLR test set | MADA | Total NLRs | Prevalence (%) |
|---|---|---|---|
| Angiosperm atlas | 6593 | 91,291 | 7.2 |
| Non-flowering plants | 34 | 2,371 | 1.4 |

**Figure S2. MAEPL motif occurrence across major plant lineages**

**A)** MAEPL motif occurrence in NLRs relative to other (non-NLR) proteins as identified through HMM profiling (hidden Markov model) representative green lineage organisms (proteomes) used throughout this study.

**(B)** MAEPL motif occurrence in NLRs relative to other (non-NLR) proteins as identified through HMM profiling (hidden Markov model) in the proteomes of model angiosperms (*Arabidopsis thaliana* and *Solanum lycopersicum*).

**(C)** MAEPL motif prevalence in non-flowering plants relative to angiosperms. An extensive set of angiosperm NLRs were queried using the angiosperm NLR atlas (ANNA).

**(D)** MADA motif prevalence in non-flowering plants relative to angiosperms. An extensive set of angiosperm NLRs were queried using the angiosperm NLR atlas (ANNA).
