## Supplementary Datasets for "The N-terminal executioner domains of NLR immune receptors are functionally conserved across major plant lineages": FigureS3.pdf

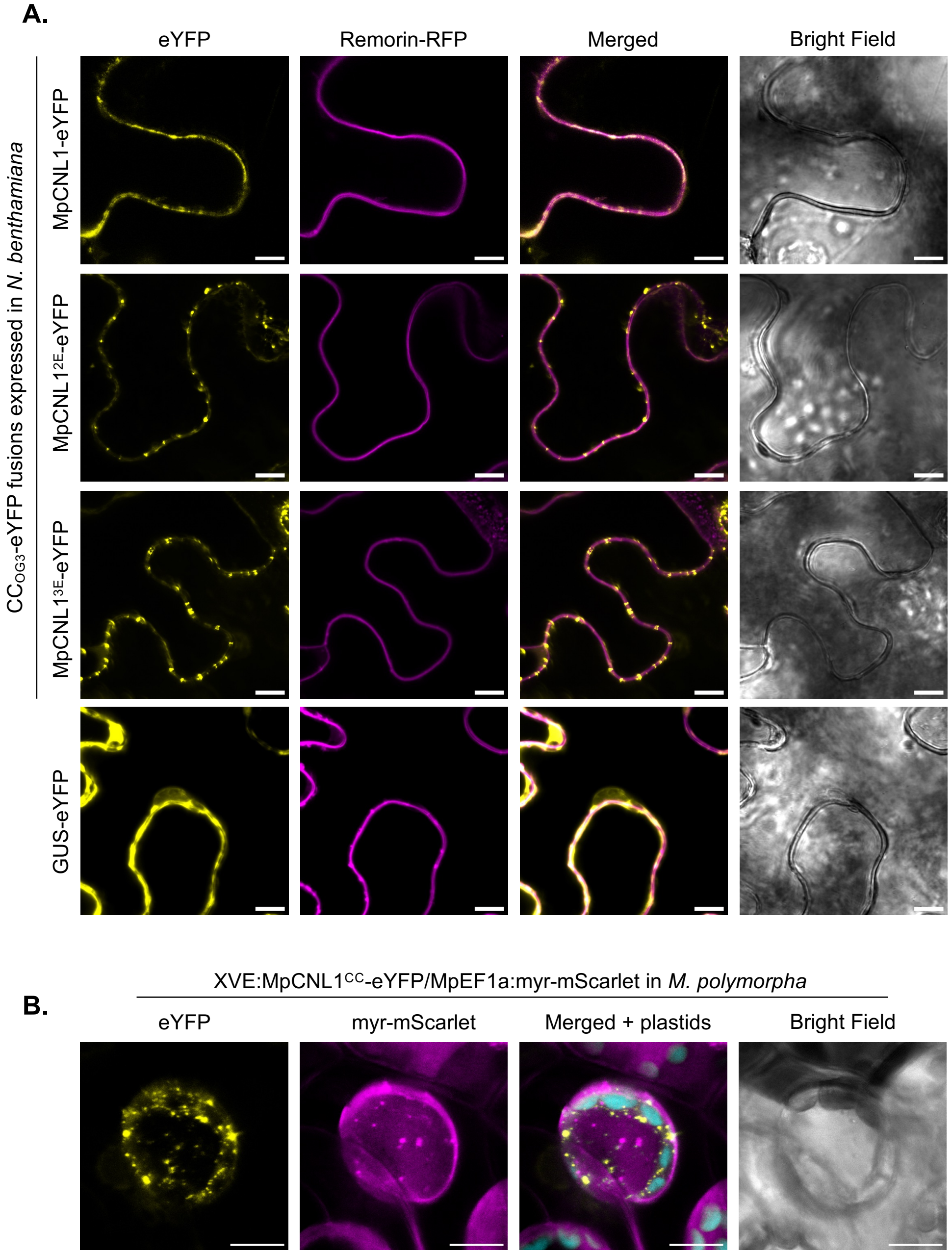


**Figure S3. Subcellular localization of MAEPL-CC$_{OG3}$ in *Nicotiana* and *Marchantia***
(**A**) Confocal fluorescence microscopy demonstrating localization of MpCNL1$^{CC}$-eYFP fusions and MAEPL motif variants (2E = L16/17E; 3E = L4/16/17E) and a GUS-eYFP control in *N. benthamiana* leaves. The REM1.3-RFP construct was co-infiltrated in combination with each construct to label the plasma membrane. Images were obtained approximately 24 hours post agroinfiltration of *N. benthamiana* leaves. Scale bars = 10 µm. Images are representative of 3 experimental replicates.
(**B**) Confocal fluorescence microscopy (Z-stack projection) demonstrating localization of MpCNL1$^{CC}$-eYFP (MpC1) alongside a myristolated-mScarlet (membrane marker) in the *Marchantia polymorpha* XVE:MpCNL1$^{CC}$-eYFP/MpEF1a:myr-mScarlet transgenic. Images were acquired 24 hours post estradiol treatment (20 µM) in liverwort thalli. Plastid autofluorescence is false-colored in cyan. Scale bars = 10 µm. Images are representative of 3 experimental replicates.
