## Supplementary Datasets for "The N-terminal executioner domains of NLR immune receptors are functionally conserved across major plant lineages": FigureS4.pdf

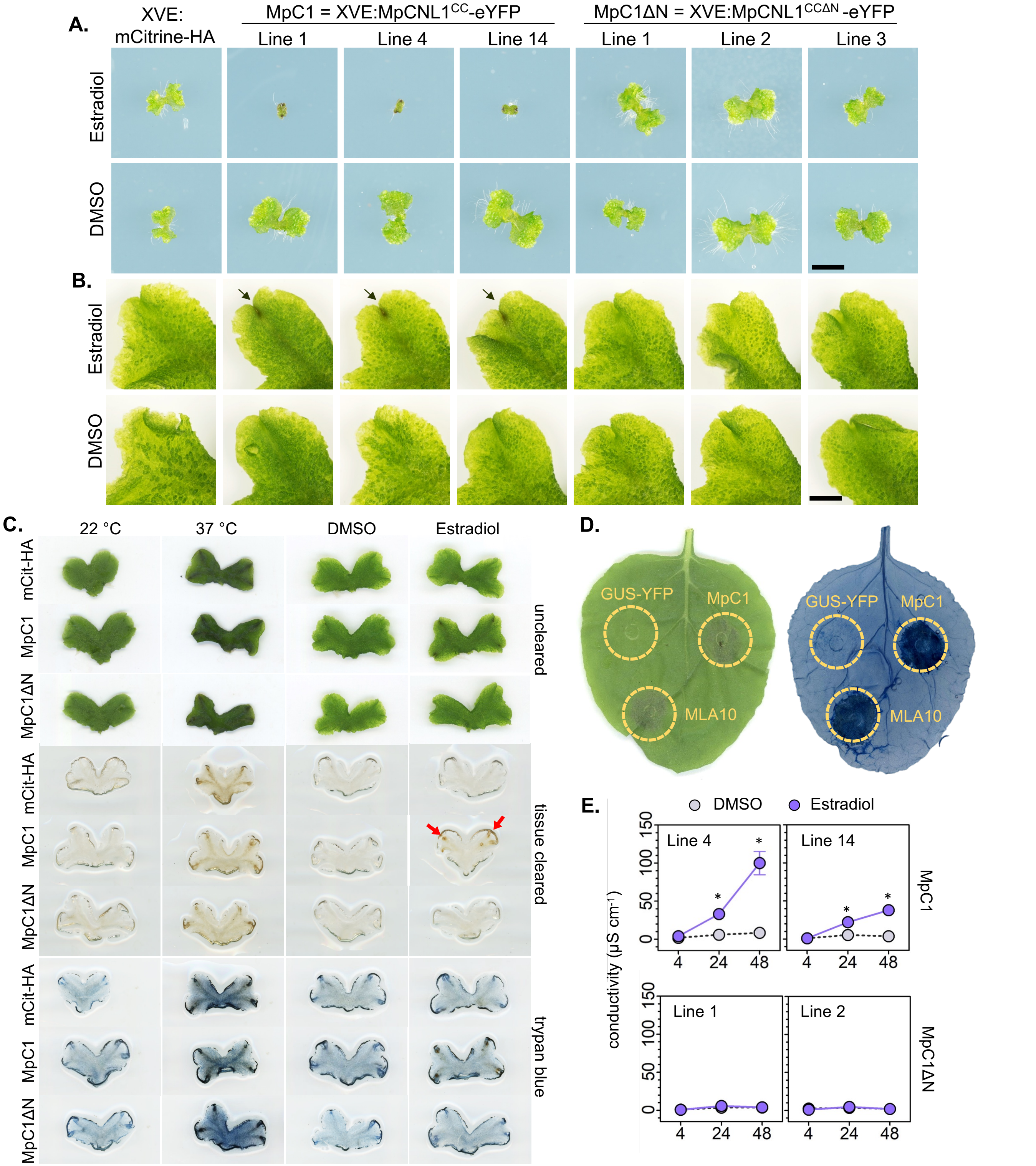


**Figure S4. Characterization of the MAEPL-CC$_{OG3}$ response in *Marchantia***

**(A)** Macroscopic phenotypes of *Marchantia* transgenic lines XVE:mCitrine-HA (mCit-HA), XVE:MpCNL1$^{CC}$-eYFP (MpC1 lines 1, 4, and 14), or the N-terminally truncated XVE:MpCNL1$^{CC\Delta N}$-eYFP (MpC1ΔN lines 1, 2, and 3) grown with estradiol (20 µM) or DMSO (0.1%) control media. Images are representative of phenotypes observed in 3 experimental replicates (n= 8 plants) at 4 days post plating. Scale bar = 25 mm.

**(B)** Macroscopic phenotypes of *Marchantia* transgenic lines (listed above) at 1 day post infiltration with estradiol (50 µM) or a DMSO control (0.25% in water). Images are representative of phenotypes observed in 3 experimental replicates (n= 8 plants). An arrow indicates tissue darkening at the apical notch of MpC1 liverworts. Scale bar = 25 mm.

**(C)** Macroscopic phenotypes and trypan blue staining of Marchantia thalli (mCit-HA, MpC1-1, MpC1ΔN-3) subjected to heat-stress (37 °C) or estradiol induction (50 µM). DMSO or 22 °C treatments were included as negative controls. Images display phenotypes 1 day post treatment in uncleared, tissue cleared (chloral hydrate), and trypan blue stained thalli. Images are representative of 3 replicates. Arrows indicate tissue darkening upon estradiol treatment.

**(D)** Cell death phenotypes (Macroscopic HR and trypan blue staining) 2 days post agroinfiltration of *N. benthamiana* leaves with MpCNL1$^{CC}$-eYFP (MpC1) and MLA10$^{CC}$-eYFP (MLA10). GUS-YFP is included as a negative control.

**(E)** Conductivity (µS cm$^{-1}$) of *Marchantia* thalli treated with estradiol (50 µM) or DMSO (0.25%) at 4, 24, and 48 hours post infiltration (hpi). Statistically significant differences are denoted by an asterisk (* $p < 0.01$, Student's t-test). Error bars represent standard deviation of the mean. Data from three independent experimental replicates is presented (n=12 plants per experiment).
