## Supplementary Datasets for "The N-terminal executioner domains of NLR immune receptors are functionally conserved across major plant lineages": FigureS5.pdf

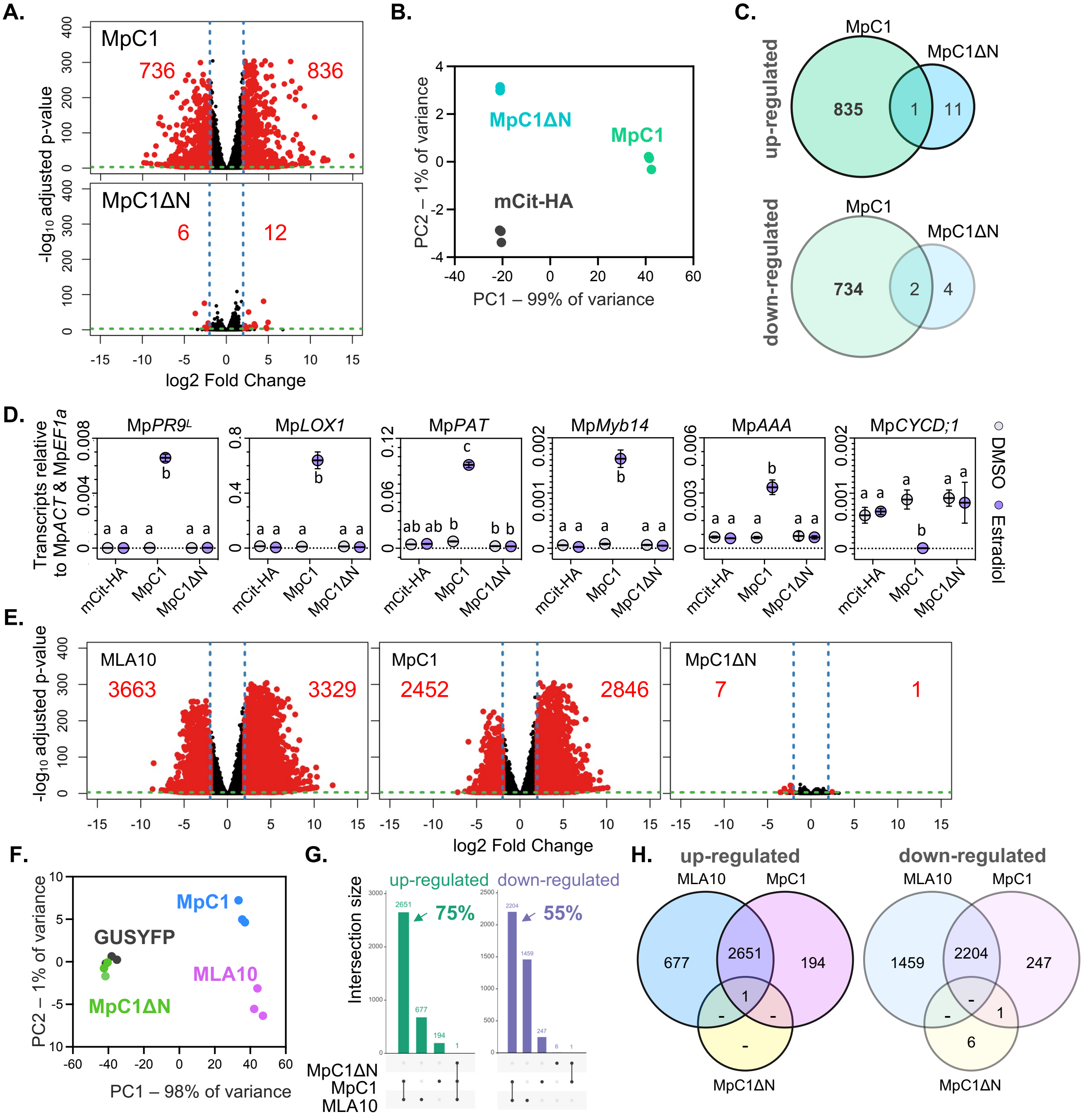


**Figure S5. Transcriptomic analysis of CC$_{OG3}$ activity in *Marchantia* and *Nicotiana***

**(A)** Volcano plots displaying pairwise differential expression analysis of estradiol-treated (50 µM) MpC1 (XVE:MpCNL1$^{CC}$-eYFP line1) or MpC1ΔN (XVE:MpCNL1$^{CC\Delta N}$-eYFP line 3) versus mCit-HA (XVE:mCitrine-HA) *Marchantia* thalli 24 hours post infiltration. Red dots represent significantly differentially expressed genes. **(B)** Principal component analysis (PCA) of *Marchantia* transcriptome replicates. **(C)** Venn diagrams depicting shared/specific differentially expressed genes (DEGs) between treatments in *Marchantia.* **(D)** qRT-PCR validation of CC marker genes in *Marchantia.* Mp*PR9$^L$*, Pathogenesis-related 9$^L$ Mp5g21510; Mp*LOX1*, Lipoxygenase Mp2g00660; Mp*PAT*, *Patatin-like phospholipase* Mp5g21760; Mp*Myb14*, R2R3 Myb transcription factor Mp5g19050; Mp*AAA*, Cell death-related AAA-ATPase Mp7g18570; Mp*CYCD;1*, D-type cyclin Mp8g17230) 24 hours after infiltration with DMSO (0.1%) or estradiol (20 µM). Expression values are displayed relative to internal Mp*ACT* and Mp*EF1a* controls. Different letters signify statistically significant differences in transcript abundance (ANOVA, Tukey's HSD, $p < 0.05$). Values represent the mean +/- standard deviation of three biological replicates ($n = 6$). This experiment was performed three times with similar results. **(E)** Volcano plots displaying pairwise differential expression analysis of MpC1 (MpCNL1$^{CC}$-eYFP), MpC1ΔN (MpCNL1$^{CC\Delta N}$-eYFP), or MLA10 (MLA10$^{CC}$-eYFP) versus the GUS-YFP control 24 hours post agro-infiltration in *N. benthamiana* leaves. Red dots represent significantly differentially expressed genes. **(F)** Principal component analysis (PCA) of *Nicotiana* transcriptome replicates. **(G)** UpSet plots showing shared and treatment-specific differentially regulated genes in *N. benthamiana*. Arrows indicate the percentage of DEGs shared between MpC1 and MLA10 treatments. **(H)** Venn diagrams depicting overlap in differentially expressed genes (DEGs) between treatments in *N. benthamiana.*
